# Automated and accurate segmentation of leaf venation networks via deep learning

**DOI:** 10.1101/2020.07.19.206631

**Authors:** H. Xu, B. Blonder, M. Jodra, Y. Malhi, M.D. Fricker

## Abstract

- Leaf vein network geometry can predict levels of resource transport, defence, and mechanical support that operate at different spatial scales. However, it is challenging to quantify network architecture across scales, due to the difficulties both in segmenting networks from images, and in extracting multi-scale statistics from subsequent network graph representations.
- Here we develop deep learning algorithms using convolutional neural networks (CNNs) to automatically segment leaf vein networks. Thirty-eight CNNs were trained on subsets of manually-defined ground-truth regions from >700 leaves representing 50 southeast Asian plant families. Ensembles of 6 independently trained CNNs were used to segment networks from larger leaf regions (~100 mm^2^). Segmented networks were analysed using hierarchical loop decomposition to extract a range of statistics describing scale transitions in vein and areole geometry.
- The CNN approach gave a precision-recall harmonic mean of 94.5% ± 6%, outperforming other current network extraction methods, and accurately described the widths, angles, and connectivity of veins. Multi-scale statistics then enabled identification of previously-undescribed variation in network architecture across species.
- We provide a LeafVeinCNN software package to enable multi-scale quantification of leaf vein networks, facilitating comparison across species and exploration of the functional significance of different leaf vein architectures.

## Introduction

Plant leaves are structured by a vein network that ranges in geometry from dendritic fans with few branches and no loops (e.g. *Ginkgo biloba*) to hierarchical forms with many loops (e.g. *Acer saccharum*) (Trivett & Pigg, 1996; Roth-Nebelsick *et al.*, 2001). The vein network has multiple roles that include transport of water, nutrients and sugars through the xylem and phloem tissues (Brodribb *et al.*, 2010; Carvalho *et al.*, 2018; Katifori, 2018), controlled deformation during bud burst (Niklas, 1999), and mechanical support and resistance to damage in the mature leaf (Sack & Scoffoni, 2013; Sharon & Sahaf, 2018). Measurements of vein architecture have provided insights into leaf development (Kang & Dengler, 2004) and evolution (Boyce *et al.*, 2009; Brodribb & Feild, 2010), prediction of leaf carbon and water fluxes (Sack & Frole, 2006; Brodribb *et al.*, 2007; Brodribb *et al.*, 2010), prediction of environmental and stress tolerances of species (de Boer *et al.*, 2012; Blonder & Enquist, 2014; Brodribb *et al.*, 2016), and reconstruction of paleo-environments from leaf fossils (Manze, 1967; Uhl & Mosbrugger, 1999; Blonder *et al.*, 2014). Venation networks are also relevant to test theories of optimal branching structure and transportation efficiency for different network architectures (Pelletier & Turcotte, 2000; Price *et al.*, 2012; Price *et al.*, 2014; Price & Weitz, 2014), particularly in response to fluctuating loads or robustness to damage (Dodds, 2010; Katifori *et al.*, 2010; Katifori, 2018).

Prediction of function is based on statistics estimated from images of vein networks (Roth-Nebelsick *et al.*, 2001). Networks have veins of different orders, varying from primary veins (attached to the petiole), to secondary (attached to the primary veins), and continuing until the ultimate veins are reached. The vein density (vein length per unit area) at each order, or across all orders, is therefore a key statistic (Uhl & Mosbrugger, 1999; Sack & Scoffoni, 2013). Measurement of the distribution of vein radii is required to determine the vein order (Price *et al.*, 2012), which is itself challenging. Alternatively vein orders can be extracted using hierarchical vein classification methods (Gan *et al.*, 2019). If vein radii can be extracted reliably, they can also be used to estimate network construction costs (John *et al.*, 2017), or to predict maximum fluxes and resource flow rates (Brodribb *et al.*, 2007; McKown *et al.*, 2010). Many networks also contain loops that partition the lamina into increasingly small areoles. For higher vein orders, qualitative terms to describe patterning, such as craspedodromous or camptodromous, have traditionally been used (Hickey, 1979; Ellis *et al.*, 2009). Quantitatively, the number of areoles per unit area provides a statistic for the amount of ‘loopiness’ (Blonder *et al.*, 2011). More detailed analysis of the nesting of loops within higher-order loops can be achieved using hierarchical loop decomposition (HLD), which constructs a binary branching tree representing the order in which adjacent areoles become connected as the intervening veins are removed (Katifori *et al.*, 2010; Katifori & Magnasco, 2012; Ronellenfitsch *et al.*, 2015; Katifori, 2018). HLD provides a range of metrics such as the Horton-Strahler index, bifurcation ratio and subtree asymmetry by analogy to river network analysis (Katifori & Magnasco, 2012; Mileyko *et al.*, 2012; Ronellenfitsch *et al.*, 2015; Katifori, 2018). Some venation networks contain freely-ending veins (FEVs) that do not form anastomoses with other veins, but which may help with efficient supply of water (Fiorin *et al.*, 2016), or possibly transport of sugars (Carvalho *et al.*, 2018). As such, density estimates of branching end-points or areole perimeter/area ratios (Kang & Dengler, 2004; Blonder *et al.*, 2018) are also useful. Finally, , the distribution of branching angles and radius or length ratios at vein branching points (Bohn *et al.*, 2002), feed into theories and models of flow efficiency, network development and architecture (Pelletier & Turcotte, 2000; Price *et al.*, 2012; Price *et al.*, 2014).

Calculating these metrics requires accurate segmentation of the venation network over a range of scales. The finest veins are typically around 10-20 μm diameter, whilst the largest veins can reach mm width in a lamina up to m^2^ in area. To capture the fine veins, networks are typically imaged after chemical clearing and staining using light microscopy or flat-bed scanning (Pérez-Harguindeguy *et al.*, 2013). X-ray imaging can also achieve high resolution with no tissue preparation, but requires access to suitable equipment (Wing, 1992; Blonder *et al.*, 2012; Schneider *et al.*, 2018). Regardless of the imaging method, accurate segmentation of the network from the image has been challenging. Images often have limited or uneven contrast, or contain unavoidable artifacts caused by other tissues and cell types, such as trichomes or glands. Furthermore, field material often includes damaged regions. As a result, simple intensity-thresholding is rarely adequate to yield useful segmentations. Several computer programs have addressed this challenge, typically using local or global enhancement filters tuned to pick out vein-like structures, either with or without human supervision. These include *phenoVein* (Bühler *et al.*, 2015), *NET* (Lasser & Katifori, 2017), *LeafGUI* (Price *et al.*, 2011), *Limani* (Dhondt *et al.*, 2012), as well as other studies that have not provided code (see for example, Bohn *et al.*, 2002; Mounsef & Karam, 2012; Larese *et al.*, 2014; Parsons-Wingerter *et al.*, 2014; Gan *et al.*, 2019). Other packages include machine learning methods to perform segmentation, e.g. *NEFI* (Rother *et al.*, 2004; Dirnberger *et al.*, 2015; Grinblat *et al.*, 2016), although these require extensive user interaction to iteratively define and correct foreground and background partitioning, and have only been applied to simple leaf vein networks (Carvalho *et al.*, 2017).

However, in our experience, none of these segmentation algorithms is robust in that it can be accurately applied across a wide range of species and image qualities, particularly for field-collected specimens, without considerable expert human guidance. Most methods are either highly tuned for specific applications, e.g. *Arabidopsis thaliana* (Bühler *et al.*, 2015), or require adjustment of many parameters produce acceptable output (e.g. Price *et al.*, 2011). Even in best-case scenarios, when these algorithms correctly classify the majority of pixels, they may fail to accurately reconstruct the topology of the network, falsely connecting or disconnecting veins, or producing incorrect vein radii. The difficulties with automated segmentation are well recognized with most studies preferring hand-traced images (e.g. Rolland-Lagan *et al.*, 2009). Indeed, current methods handbooks only recommend expert hand-tracing to estimate venation network properties (Pérez-Harguindeguy *et al.*, 2013). Nevertheless, while hand-tracing is accurate, it is slow and time-intensive, limiting throughput of samples and restricting the area of leaf analyzed.

Here we develop an alternative deep learning method to segment minor vein networks from high-resolution cleared leaf images. The approach is motivated by our observation that humans can trace venation networks consistently and accurately after minimal training, despite low contrast and many artifacts in the underlying images. We therefore describe an approach for training an ensemble of Convolutional Neural Networks (CNNs) to learn how to carry out this segmentation task based on ground-truth (GT) tracings previously made by humans from field-collected leaves of tropical rainforest trees. The trained CNNs are provided within the LeafVeinCNN software package which can be applied to any high-resolution cleared leaf image.

## Materials and Methods

### Leaf vein imaging

A set of calibration leaves were sampled from eight permanent forest plots in Sabah state, Borneo, Malaysia (Riutta *et al.*, 2018), which are characterized by a mixed dipterocarp lowland forest. A total of 727 samples were obtained from 295 species (50 families). The sampling protocols and non-vein datasets have been described previously (Both *et al.*, 2018), and the complete set of vein images published (Blonder *et al.*, 2019).

A 1 cm^2^ sample was cut from the middle of the lamina, excluding any primary veins, chemically cleared, stained, and slide-mounted following standard protocols (Pérez-Harguindeguy *et al.*, 2013; Blonder *et al.*, 2018). Each sample was imaged using a compound microscope (Olympus, BX43) with 2x apochromat objective and an Olympus SC100 color camera (3840 × 2748 pixel resolution). 9-16 overlapping image fields were stitched together to obtain a complete image of the sampled area using the image capture software. Final images had a resolution of 595 pixels mm^−1^ and a typical extent of ~7000 × 7000 pixels.

Images were pre-processed by retaining the green channel (which had the greatest contrast), and applying contrast-limited adaptive histogram equalization (CLAHE, Zuiderveld, 1994), with a 400 × 400 pixel window and a contrast limit of 0.01.

### Ground-truth tracing

All veins within a polygonal region-of-interest (ROI) of approximately 700×700 pixels were manually traced at full-width for each image, using a digitizing tablet (Cintiq 22HD, Wacom) and image processing software (GIMP). In addition, any veins in the complete image with width >200 μm were manually traced, as these veins were not routinely included in the GT-ROIs used to train the CNNs. Inclusion of manual large veins avoided over-segmentation of texture within the larger veins which were occasionally recognized by the CNN as separate veins. Alternatively, we used down-sampled images with the same CNN to automatically segment larger veins at full width. Lastly, a separate mask was manually traced to exclude damaged or background regions from the full image.

Each tracing required ~30 min. of human time. Each image layer in GIMP (i.e. the ROI, large vein or mask) was drawn by a single person and was reviewed and subsequently edited by 2-3 other people until consensus was reached, to give a final set of 727 GT images.

### Convolutional Neural Network (CNN) segmentation using U-Net

CNNs are a deep learning method that allow automated extraction of features from images (Krizhevsky *et al.*, 2012; LeCun *et al.*, 2015). They differ from classic machine learning methods in that they synthesize information at multiple scales via a sequence of convolution, pooling, and up-sampling operations. Convolution-deconvolution networks, such as U-Net (Ronneberger *et al.*, 2015), use a set of rectangular kernels that pick up different features in the image at each scale, which are then pooled and combined in each network layer to ultimately yield pixel-level classifications. ‘Deep’ neural networks include multiple convolutional layers whilst ‘learning’ occurs via a backpropagation algorithm (Werbos, 1974) that optimises the weights of the network by comparing the output result to the GT.

We used a CNN based on the U-net architecture (Ronneberger *et al.*, 2015). The first convolutional layer comprised 32 small (3 × 3) kernels that were convolved with the image to give 32 feature maps (Fig. 1). The feature maps were batch-normalized to give a mean of zero and variance of 1 to speed up training (Ioffe & Szegedy, 2015), followed by non-linear rectification (ReLU) that sets negative inputs to zero to improve training performance (Glorot *et al.*, 2011). A second convolution using another set of 32 small (3 × 3) kernels was used to increase the receptive field in this layer (Simonyan & Zisserman, 2014), followed by batch-normalization and ReLU (Fig. 1). The size and resolution of the input feature maps and the output feature maps were preserved when passing through the convolutional blocks, with padding applied during convolutions. To generate the next layer, the output from each convolution block was subject to non-linear down-sampling using the maximum from a 2 × 2 pixel window with a stride of 2, effectively halving the resolution of the feature maps in both dimensions. This gave five levels of feature maps in the down-sampling arm of the network which provided increasingly higher-level representation of the input image patches.

**Figure 1:**
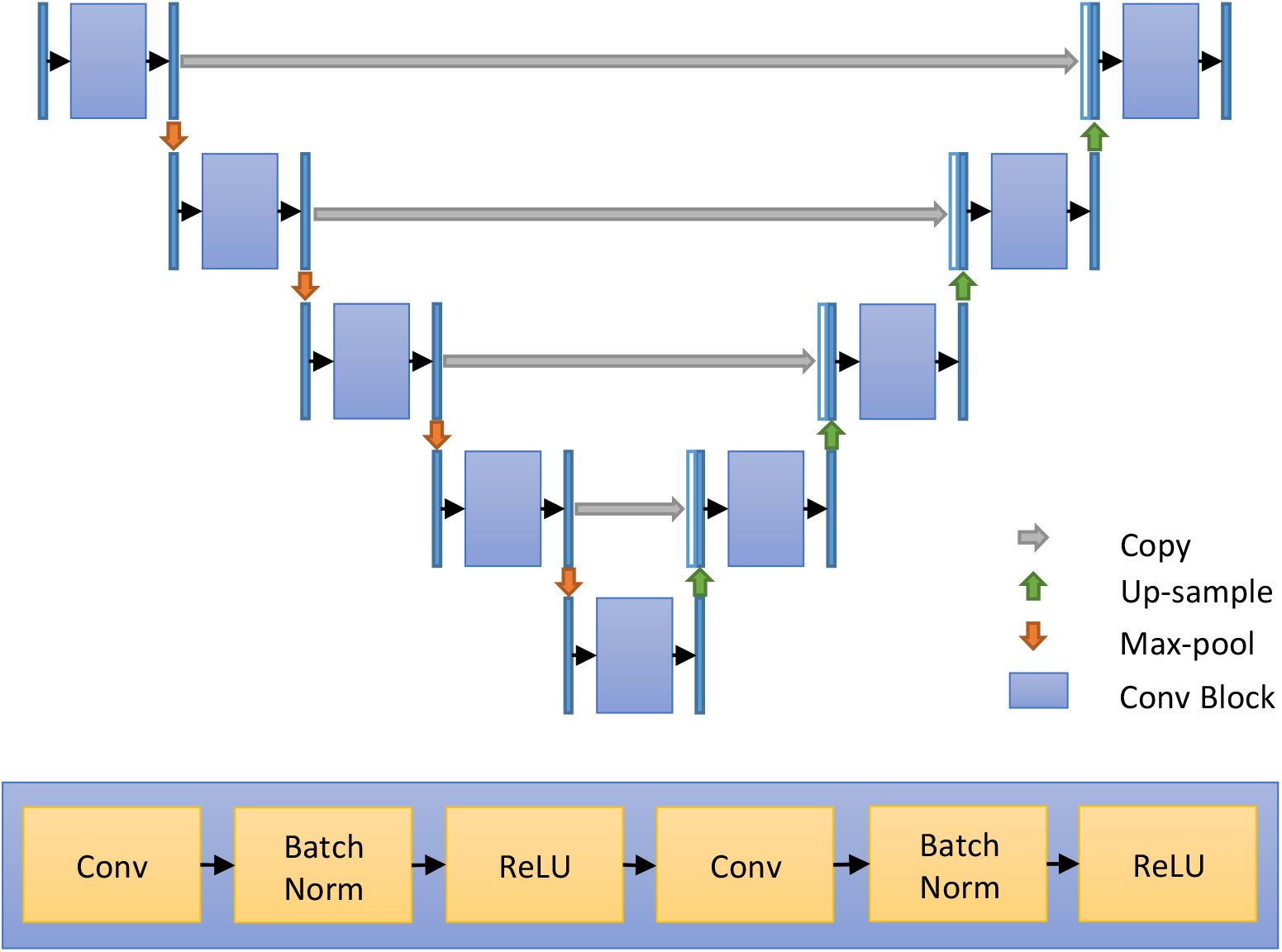
Schematic diagram of the U-Net CNN architecture. Each layer in the CNN is built of two concatenated blocks comprising a convolution layer, followed by batch-normalisation and non-linear rectification (ReLU). This is followed by 2×2 max pooling to reduce the scale in the down-sampling arm (left). In addition, a copy of the output of the convolution block is appended to the up-sampling arm (right) at the same level that retain the high-resolution information required to yield per-pixel outputs. The final layer is fully connected and provides a probability classification into vein or non-vein pixels using a sigmoid function.

We also followed the approach of U-Net and Fully Convolutional Networks (FCN) (Long *et al.*, 2015), and kept the same number of feature maps for layers with the same resolution, whilst doubling the number of feature maps when the resolution was halved by down-sampling, and halving the number of feature maps when resolution was doubled in the up-sampling arm. The feature maps at each down-sampled layer were also copied and concatenated with those at the same resolution in the up-sampling arm, to give feed-forward shortcut connections that retain the high-resolution information required to yield a per-pixel output (Long *et al.*, 2015). The activation function of the final fully connected output layer was sigmoid and gave a 1:1 pixel classification of the vein probability. This resulted in nine convolutional blocks in total (Fig. 1).

### Data augmentation and training data sets

The manually digitized GT images were used to train and evaluate the CNN. Each GT region was resampled to augment the training data and to prevent over-fitting. 32 samples per image, fully contained within the GT region, were extracted with randomized variation in the box size (256 × 256 ± 20% pixels), rotation angle (between 0 and 2π radians), (*x,y*) shift in centroid coordinates, and randomized image reflection. Augmented samples were then resized to 256 × 256 pixels (Fig. 2).

**Figure 2:**
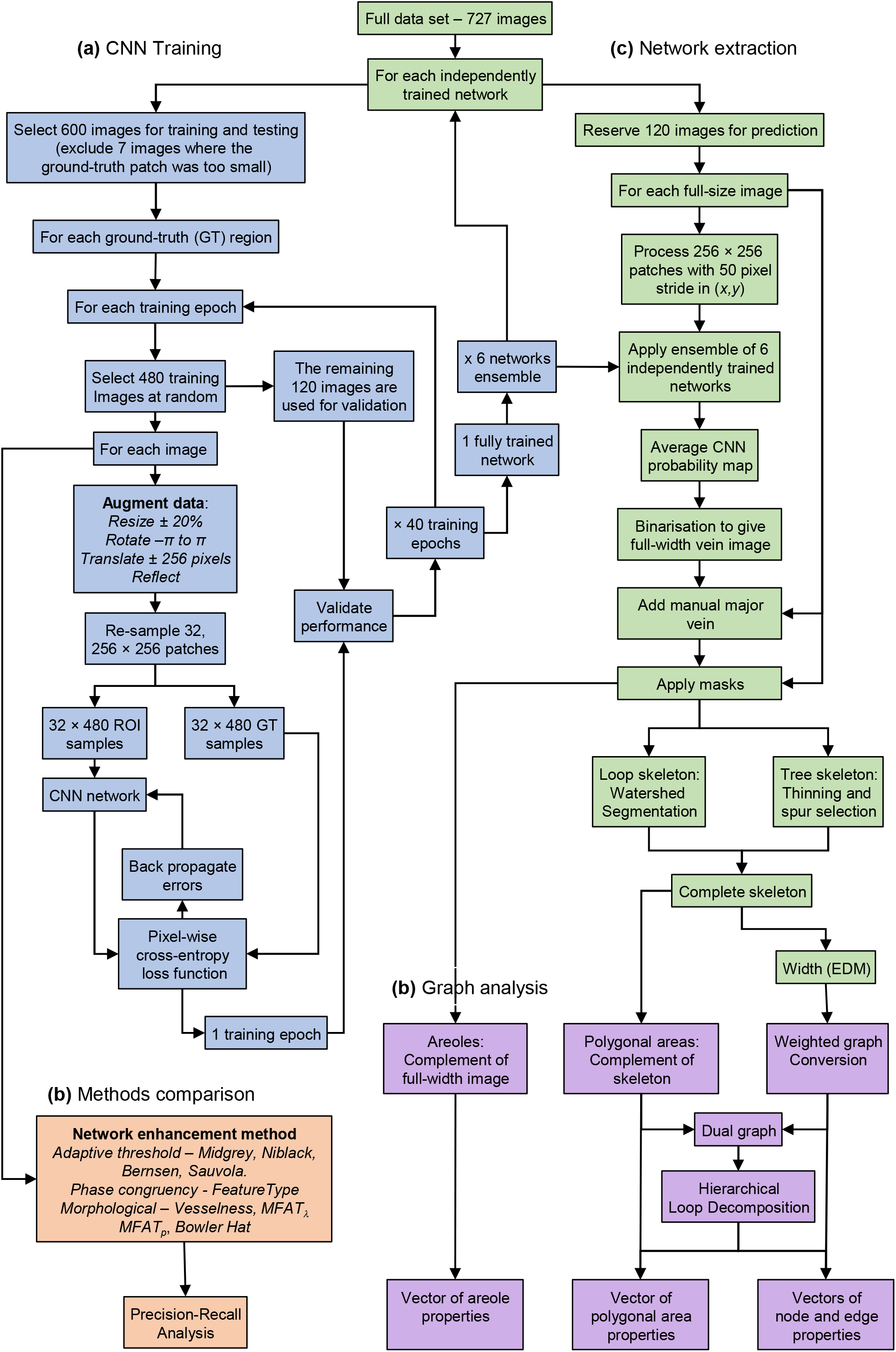
Flow diagram of the major steps in CNN training, segmentation and analysis. At each iteration, one set of 120 non-overlapping images is reserved for prediction and network extraction (green boxes). At each training epoch (blue boxes), 480 images are selected at random for training from the remainder, and a further 120 reserved for validation **(a)**. For each training image, a region-of-interest (ROI) and the associated manually delineated full-width ground-truth (GT) are augmented by scaling, translation, rotation and reflection to give 32, 256×256 image patches. The patches are fed to the CNN, and the output compared to the augmented GT patches using the Dice Similarity Coefficient (*F*_*1*_ statistic). The errors are back-propagated through the network to adjust the internal weights, and the process repeated with another training image. Once the network has been trained on all 480 initial images, termed an epoch, the output is tested against the validation set. Training continues for up to 40 epochs to give a fully trained network, and then repeated to give an ensemble of six independently trained networks. These are used to predict the vein architecture in the set of reserved test images. Complete prediction for the complete data set requires seven iterations. (**b**) The performance of the CNN approach is compared to a range of different network enhancement and segmentation methods (orange boxes), including local adaptive thresholding (Midgrey, Niblack, Bernsen and Sauvola), or multi-scale ridge enhancement methods (Vesselness, MFAT_λ_, MFAT_p_, Bowler Hat and Phase congruency), using Precision-Recall (PR) analysis. (**c**) To extract the vein network, the full-size images are processed through the ensemble of independently trained CNN networks to give an average probability map of the vein network (green boxes). The probability map is segmented to give a full-width binary image (FWB) and then skeletonized in a two-step process to give loops and trees. The width of the veins is estimated from the Euclidean Distance Map (EDM) of the FWB image. (**d**) The skeleton is converted to a weighted graph representation with nodes at junctions linked by edges along the veins, which are both associated with a vector of properties (purple boxes). The areoles are extracted from the complement of the FWB image, whilst the polygonal regions, including the area of the veins up to the skeleton are extracted from the complement of the skeleton. In addition, a weighted dual-graph is constructed that links adjacent areoles (nodes) with edges weighted by the width of the intervening vein. Edges are removed in sequence to fuse the parent areoles and reveal the nested loop structure of the network by Hierarchical Loop Decomposition (HLD). Network metrics are calculated at each step of the HLD to track the vein architecture at different scales.

### Network Training

The CNN was initialized using the Glorot uniform initializer with random weights drawn from a uniform distribution (Glorot & Bengio, 2010). At the start of each training run, 120 non-overlapping images were reserved as the test data set. For each training epoch, 480 leaf images were randomly selected from the remaining images and the manually digitized ROIs used as training samples. The remaining 120 images were used to validate the network performance. The trained CNN was then used to segment the set of 120 full-size test images that had not been used in training or validation. This required seven iterations to segment all of the original full-size images. The process was repeated to give an ensemble of six independently trained networks for each image (Hansen & Salamon, 1990; Sollich & Krogh, 1996). A flow diagram of the process is shown in Fig. 2. Of the 727 samples in the dataset, seven had digitized regions which were too small for image patch extraction, which were excluded from training, but were still used for validation. At each training iteration, a mini-batch of eight 256 x 256 patches, constrained by the memory limit of each GPU, was sampled and fed to the network in sequence to ensure robust convergence by avoiding local minima. The pixel-wise cross entropy was used as the loss function during training, and the Dice similarity metric was used for network selection. Networks were typically trained for 20-40 epochs, where an epoch was a complete run through one set of 480×32 samples, with a new training data set used at each epoch (giving a total of 480×32×40 augmented training samples) until the error asymptotically approached a stable value with <0.1% variation.

The U-Net algorithm was run on one of two clusters, either with 2x Intel E5-2609v2 processors (total 8 cores at 2.5 GHz, 128GB RAM), or 2x Intel E5-2609v3 processors (total 12 cores at 1.9 GHz, 128GB memory). Image processing occurred in parallel on three graphical processing units (GPUs) (nVidia, GeForce GTX 1080 Ti, each with 3580 cores and 11GB memory). All CNN code was implemented in Keras with Tensorflow using Python 3. Training the CNN on the calibration data required 8 min per epoch using 3 GPUs and 1 CPU.

The trained networks were also imported into MATLAB (Mathworks, Natick, MA) for incorporation into a GUI interface (see user manual and software provided in Supplementary Information). This required replacement of the Keras *Upsampling2D* layers with MATLAB *transposedConv2dLayer* layers using a stride of 2 and uniform weights equal to 1.

### Vein prediction on the full-size images

For vein prediction over the full 10 mm × 10 mm region, images were processed as overlapping 256 × 256 pixel patches. At each iteration, a mini-batch of eight patches was sampled at random without replacement, with a separation (stride) of 50 pixels in both *x* and *y* to give >25-fold over-sampling, and a small overlap between patches. The results were assembled back to the original image size to give a probability map of predicted values, with each pixel classified at least 25 times within different image contexts due to the stride overlap. Calculation of the CNN on a full-sized image took ~8 minutes on the GPU clusters. The analysis code running on a single CPU took ~20 min depending on the complexity of the network and amount of down-sampling. The typical processing speed was approximately 60,000 px s^−1^.

### Conversion of the CNN probability map to a weighted network

For each full-size leaf image (Fig. 3a), the six CNN probability maps were averaged to generate a mean probability map (Fig. 3b), down-sampled by a factor of 2 to reduce subsequent computational costs, and binarised using a threshold based on the average *F_β_* statistic from Precision-Recall analysis (see below). Only the largest connected component was retained. Damaged and background regions that were defined manually for each image were masked from analysis.

**Figure 3:**
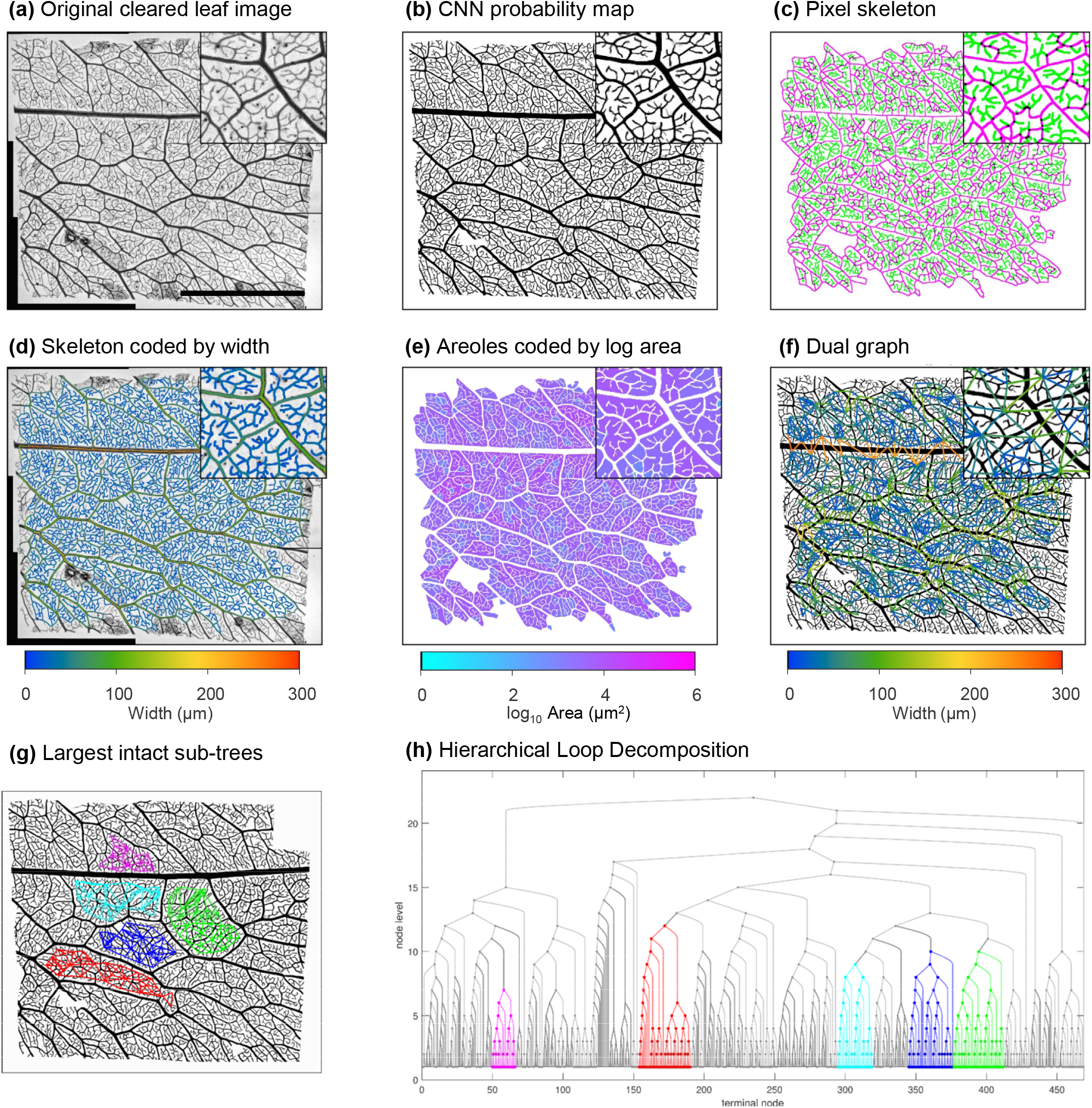
Processing steps in CNN training, segmentation and analysis. (**a**) Original image of a typical cleared leaf sample following CLAHE enhancement, with inset showing the central region at 6x zoom. Scale bar = 5 mm. (**b**) Average ensemble CNN probability map. (**c**) Pixel-skeleton with loops in magenta and trees in green. (**d**) Pseudo-colour coded vein width skeleton superimposed on the original image. (**e**) Areoles, pseudo-colour coded by log10 area. (**f**) Dual graph connecting centroids of adjacent areoles with the edge weight given by the average width of the intervening vein. (**g**) Overlay of the dual graph for the 5 largest intact subtrees in the full-size image. (**h**) The complete HLD of the dual-graph including the five largest intact sub-trees following the colour scheme in (**g**).

The resultant full-width binary (FWB) image was thinned to a single pixel wide skeleton (Fig. S1b). Simple thinning algorithms gave an irregular zig-zag backbone around junctions. A better centerline was realized using a hybrid skeletonisation protocol that combined watershed segmentation to find all closed loops (Figure **3c**, magenta), with thinning to extract tree-like branches (Figure **3c**, green). In brief, watershed segmentation was applied to the Euclidean Distance Map (EDM) from the FWB image to find all closed loops (Fig. 3c, magenta). Tree-like branches were extracted as the difference between the complete thinned skeleton (Zhang & Suen, 1984), and the skeleton after removal of spurs (Fig. 3c, green). The ‘trunks’ of the tree-like skeleton were then re-connected to the nearest pixel in the watershed loop skeleton, determined from the nearest-neighbor transform of the EDM, and the pixel positions written back into the image using the Bresenham line algorithm (Bresenham, 1965). This hybrid approach had the benefit that loops and tree-like portions of the skeleton were automatically separated and could be analysed separately (Fig. 3c).

The pixel skeleton was converted into separate vein segments using the Matlab *edgelink* algorithm (Kovesi, 2000), which provided a set of pixel-coordinates for each vein in the network. The vein segments were converted to a graph representation (Fig. S1c) with nodes (N) assigned to each junction or free end connected by edges (E) (Roth-Nebelsick *et al.*, 2001).

The width at each pixel of the skeleton was calculated from the EDM. Pixel width estimates were averaged to give an initial width estimate for each vein. However, this over-estimated the width for small veins branching from larger veins, as it included values where the smaller vein overlapped the larger one (Bohn *et al.*, 2002). Thus, the width for the smallest vein at each junction was refined by excluding pixels in the overlap region, before taking the average of the remaining pixels.

To visualise the segmented network, the average center-width of each edge was mapped back into the pixel skeleton, colour-coded on a perceptually uniform rainbow scale, and superimposed on the original leaf image (Fig. 3d). The areole regions between the veins (i.e. excluding the veins themselves) and the area of the polygonal region up to the pixel skeleton were determined from the complement of the FWB image, or the skeleton, respectively (Fig. 3e).

### Single scale network metrics

Multiple metrics were extracted from the graph representations for each vein (Fig. S1d, Table S2), node (Fig. S1e, Table S3), areole, and polygonal area (Fig. S1f, Table S4 and S5, respectively). In addition, a set of summary statistics describing the distribution of each metric were also calculated (Table S7), adopting a comparable terminology to Larese *et al.,* (Larese *et al.*, 2014), with prefixes *V* - veins, *N* - nodes, *A* - areoles, and *P* - polygonal areas. Vein metrics were further sub-divided into the total vein network (*Tot*), veins forming loops (*Loop*), and veins forming trees (*Tree*). The summary statistics were drawn from mean (*av*), median (*md*), min (*mn*), max (*mx*), standard deviation (*sd*), or skewness (*sk*) of each metric distribution. In the case of angular metrics, values were summarized using circular statistics (Berens, 2009). Each summary statistic was then given a three-letter code.

The revised ‘center-weighted’ width value (three-letter code, *Wid*), along with the corresponding length (*Len*), excluding vein overlap and measured as the sum of the Euclidean distance between edge pixels, surface area (*SAr*), volume (*Vol*), tortuosity (*Tor*), and orientation (*Ori*) were added to the vector of edge properties, along with the average edge intensity (*Int*) and average CNN probability (*Prb*) (Figure S1d). A vector of properties was also calculated for each node that included the node degree (*K*), diameter of the parent branch (*BD0*), and daughter branches (*BD1* and *BD2*), their orientation (*OD0, OD1,* and *OD2*), and branching angles (*A10, A20,* and *A21*). The angular metrics were calculated for straight-line segments joining the node and the midpoint of each vein (Figure S1e)

### Areole and polygonal area metrics

A range of morphological features were automatically calculated for each areole (*A*) and polygonal region (*P*) (Figure S1f, Supplementary Table 3 and 4, respectively), along with summary statistics labelled with a three letter code, including the area (*Are*), convex area (*CnA*), eccentricity (*Ecc*), equivalent diameter (*EqD*), perimeter (*Per*), major (*Maj*) and minor (*Min*) axes, orientation (*Ori*), solidity (*Sld = Are*⁄*CnA*), elongation *(Elg = Maj*⁄*Min)*, circularity (*Cir = 4 × π × Are*⁄*Per*^*2*^) and roughness (*Rgℎ = Per*^*2*^⁄*Are*), maximum (*Dmx*) and mean (*Dav*) distance to the nearest vein.

### Dual-graph representation and hierarchical loop decomposition (HLD)

A dual-graph of the vein skeleton was calculated that joined adjacent polygons, with nodes placed at the centroid of the polygons, and the connecting edges weighted using the width of the intervening vein segment (Fig. 3f). The dual-graph was converted to a nested branching tree using hierarchical loop decomposition (HLD, Katifori & Magnasco, 2012) by progressively removing the thinnest edges in sequence and fusing the adjacent polygons until only a single polygon remained (Fig. 3g,h). This process determined how vein loops are nested within larger vein loops across spatial scales. A subset of vein and polygonal area metrics were calculated at each step, allowing analysis of network metrics across the scale of the fusion events (SI Table 6). Analysis was restricted to fully bounded regions (Fig. 3g) to ensure metrics were not influenced by incomplete veins or areoles truncated at the boundary, although this also restricted the number of scales available in each sample.

### Precision-Recall analysis

The performance of the CNN classification, and selection of the optimum threshold, were determined by Precision-Recall (P-R) analysis. The number of true positives (*T*_*P*_), true negatives (*T*_*N*_), false positives (*F*_*P*_) and false negatives (*F*_*N*_) were calculated against the manually-defined GT image. In the case of comparisons of skeleton extraction, a tolerance of 3 pixels (~10 μm for the down-sampled images) was allowed (Lopez-Molina *et al.*, 2013). P-R analysis was used in preference to Receiver Operating Characteristic (ROC) plots, as the former is better suited to imbalanced data sets, where *T*_*N*_ from the background is expected to be much greater than *T*_*P*_ from the skeleton (Saito & Rehmsmeier, 2015). Precision (*P)* was calculated as = 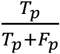, recall (*R*) as 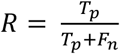 and overall performance assessed with the Dice similarity metric (*F*_*1*_) score as the harmonic mean of precision and recall, 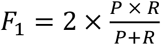 or the more generalised *Fβ* score, where 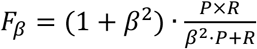, in this case using *β = 2*, which weights recall higher than precision. The threshold that gave the highest *F*_*1*_ or *F*_*β2*_ score was used to segment the image.

### Comparison with other vein segmentation methods

To compare the performance to other methods, the GT-ROI was analysed using a range of standard local adaptive thresholding methods, including Midgrey, Niblack (Niblack, 1985), Sauvola (Sauvola & Pietikäinen, 2000), and Bernsen (Bernsen, 1986) algorithms, effectively extending the local adaptive threshold approach used in *LeafGUI* (Price et al., 2011; Green et al., 2014). Each method was applied to an inverted version of the original image, with a disk shaped local neighbourhood with a radius of 45 pixels, slightly greater than the largest vein in the down-sampled GT images. The offset constant was varied from −0.5 to 0.5 (for Midgrey and Niblack) or 0 to 1 for the other algorithms, in 0.05 increments, and the optimum threshold value selected using the *F*_*1*_ or *F*_*β2*_ metric from the P-R analysis.

To emulate the multiscale Hessian-based Vesselness approach used in *phenoVein* (Bühler et al., 2015), the Matlab fibermetric implementation of the Frangi ‘Vesselness’ filter (Frangi et al., 1998) was used with vein thickness between 4-9 pixels, applied over four scales using a Gaussian image pyramid (Burt & Adelson, 1983) to cover the largest veins. We also tested improved Hessian-based enhancement techniques using Multiscale Fractional Anisotropy Tensors (MFAT), in both their eigenvalue-based (MFAT_λ_) and probability-based (MFAT_p_) form (Alhasson *et al.*, 2018), and the intensity-independent, multiscale phase-congruency enhancement developed by Kovesi (Kovesi, 1999; Kovesi, 2000), that we have previously used to segment fungal (Obara et al., 2012), slime mold (Fricker et al., 2017) and ER networks (Pain *et al.*, 2019). We used the normalized local weighted mean phase angle (‘Feature Type’) to give intensity-independent enhancement of vein structures, initially calculated over 3-5 scales and 6 orientations, and then applied to an image pyramid to cover larger scales. In both the vesselness and phase-congruency enhancement, noise was suppressed by setting all variation less than 0.1-0.2 in an extended minimum transformation to zero. As an alternative morphology-based approach, we also tested the multiscale *Bowler Hat* algorithm (Sazak *et al.*, 2018), which is designed to be more robust at junctions and less sensitive to interference from blob-like inclusions. In all cases, the resultant multiscale representation was collapsed to a single image using a maximum projection and segmented over varying thresholds from 0 to 1 in 0.05 increments to find the optimum performance as judged by the maximum *F*_*1*_ or *F*_*β2*_ statistic in Precision-Recall comparison to GT. A full set of processing parameters for these methods is given in Table S1.

As the vein enhancement approaches may be better suited to extraction of the pixel skeleton, rather than a binary image covering the full-width of the vein, the P-R analysis was also run following conversion of the binary image at each threshold value to a single-pixel wide skeleton.

### Data and algorithm availability

All MATLAB scripts and the standalone LeafVeinCNN software and manual (Fig. S2) are available from (URL provided on acceptance). The dataset, including GTs, is publicly available (Blonder *et al.*, 2019). The CNN predictions and networks for the image dataset are available from the Oxford Research Archive (ORA) (URL provided on acceptance).

## Results

### CNNs provided high accuracy vein network segmentation

To illustrate the accuracy of the ensemble CNN approach, we show weighted networks for six species (*Artocarpus odoratissimus*, *Dryobalanops lanceolata*, *Lophopetalum javanicum*, *Macaranga pearsonii*, *Pentace laxiflora*, and *Terminalia citrina*) covering a range of different vein architectures from dense loops to open trees (Fig. 4). In each case the centre-line of each vein was correctly identified, the topology of the complete network was accurately segmented, and the width corresponded to the vein thickness.

**Figure 4:**
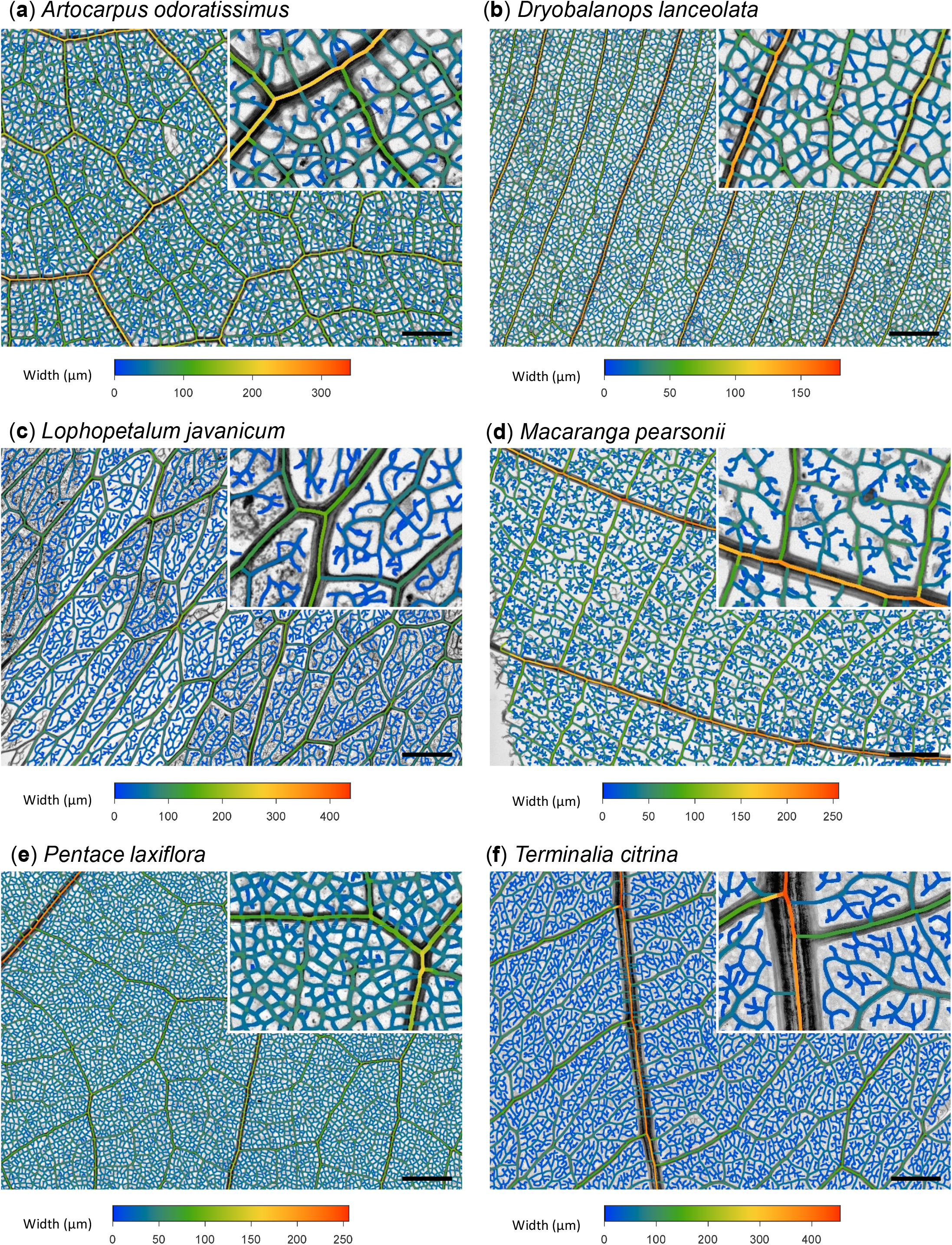
Typical CNN segmentation results. Six example leaf venation networks (grayscale images) overlaid with the extracted network skeleton pseudo-color coded to represent vein thickness on a rainbow scale (blue; thin; red, thick). Inset panels show a 3-fold zoom of center-located ROI. Scale bars = 10 mm.

To quantify ensemble CNN performance, we examined a masked ROI from each full-size image (Fig. 5a) containing the manually delineated GT (Fig. 5a’). The ensemble CNN applied to this ROI provided a smooth, high-contrast probability map of vein identity (Fig. 5b) that was thresholded to give a FWB image (Fig. 5b’) and compared to the GT using P-R analysis (Fig. 5l). The value used to threshold the CNN probability map was systematically varied to generate a P-R curve, and the optimum value determined from the maximum *F*_*1*_ or *F*_*β2*_ metrics (see Fig. 5l, circled and asterisk points, respectively). The P-R image constructed using the *F*_*β2*_ threshold value (Fig 5b’) showed that the majority of the segmentation matched the GT (green), whilst the very tips of the FEVs were slightly clipped (shown as false negatives, *F*_*N*_ in red), and the vein widths slightly over-estimated with some false positives (*F*_*P*_, shown in blue).

**Figure 5:**
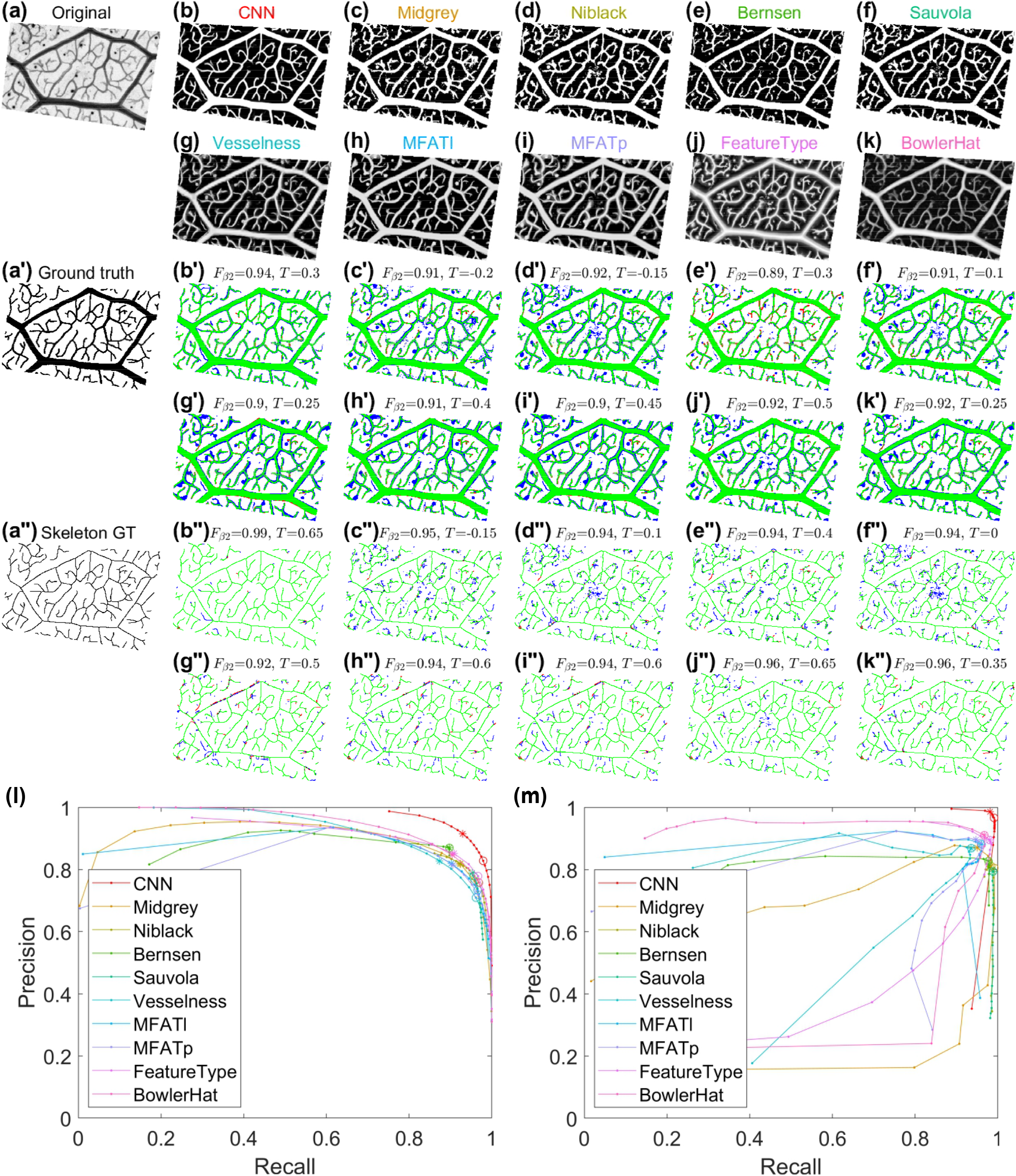
Comparison of different network enhancement and segmentation methods. A small ROI from the full-size image (**a**), was processed using different enhancement algorithms including the trained CNN (**b**); local adaptive thresholding using Midgrey (**c**), Niblack (**d**), Bernsen (**e**) or Sauvola (**f**) algorithms; Hessian-based vessel enhancement using Vesselness (**g**), MFAT_λ_ (**h**) and MFAT_p_ (**i**); Phase-congruency Feature Type (**j**), and Bowler Hat morphological filtering (**k**). The performance of each method was evaluated against a manually-defined FWB-GT image (**a’**) or the resultant single-pixel skeleton (**a”**). In each case, true positives (*T*_*P*_) are coded in green, false negatives (*F*_*N*_) in red, and false positives (*F*_*P*_) in blue for the FWB image (**b’-k’**), and the single-pixel skeleton (**b”-k”**). Precision-Recall (P-R) plots illustrate the performance of each method as the threshold is varied for the full-width comparison (**l**) and following skeletonisation (**m**). For each algorithm the optimum threshold was selected using the maximum value of the Dice Similarity Coefficient (*F*_*1*_, asterisk), or *F*_*β2*_ statistic (open circle) from the P-R plots. Values are shown above the corresponding image for the *F*_*β2*_ statistic, and the optimal threshold value T. The CNN gave the best performance for both the FWB image (*F*_*β2*_ = 0.943) and the skeleton (*F*_*β2*_ = 0.99) comparison. In addition, the P-R plots showed the CNN method performed well over a wide range of segmentation thresholds, whilst the other algorithms were extremely sensitive to the precise threshold used.

Ensemble CNN performance was affected by the contrast range present in the image initially (Fig. S3), but this could be completely recovered using CLAHE to enhance contrast (Fig. S4). Furthermore, performance was stable within a wide range of CLAHE settings (Fig. S5), suggesting the most critical feature is to cover the full contrast range, rather than the fine tuning the local adaptive features. Increasing levels of blur degraded CNN performance (Fig. S6), but only at levels that would be unacceptable out-of-focus in the original image. Possibly the strongest effect on CNN performance was the pixel resolution of the original image, where fine veins were progressively lost as the resolution decreased (Fig. S7). Nevertheless, performance was almost completely recovered by simple image interpolation back to the standard pixel size (Fig. S8).

### CNNs performed better than other network extraction algorithms

P-R curves for the other enhancement methods were worse that the CNN result (Fig. 5l). All the local adaptive thresholding approaches tested (Midgrey, Niblack, Bernsen and Sauvola, Fig. 5c-f, respectively) captured much of the main vein structures, but missed or fragmented fine veins (Fig. 5c’-f’, red), and included some noise or non-vein structures (Fig. 5c’-f’, blue). The Hessian based techniques (‘Vesselness’, MFAT_λ_ and MFAT_p_, Fig. 5g-i, respectively) showed slightly greater selectivity for the veins over background compared to adaptive thresholding methods, but had errors in the width estimate (Fig. 5g’-i’). The ‘Vesselness’ retained more of the intensity information from the original image and tended to give low intensities at junctions (Fig. 5g, g’), whilst MFAT_λ_ and MFAT_p_ gave higher contrast and better resolution of junctions (Fig. 5h-i, h’-i’). The phase-congruency Feature-Type enhancement gave high contrast and even intensity for all veins, irrespective of their size, and captured the topology quite well (Fig. 5j), but gave numerous small errors along the boundary of the binarised veins, and included discrete round structures that we infer to be glands (Fig. 5j’). The Bowler Hat enhancement (Fig. 5k) improved the selection of veins over non-veins, but was still dominated by the original intensity information, making subsequent threshold selection difficult (Fig. 5k’).

As several of these enhancement methods were originally designed to facilitate extraction of the skeleton, rather than FWB image, we also compared their performance against the GT skeleton (Fig. 5a”-k”). The performance of the CNN was even better in this comparison (*F*_*1*_ = 0.98, *F*_*β2*_ = 0.99), and was robust to threshold selection (Fig. 5m), with the optimal threshold across all species of 0.534 ± 0.202 s.d. for *F*_*1*_, 0.378 ± 0.197 s.d. for *F*_*β2*_. The other methods lagged behind the CNN result, but typically showed better P-R performance for the skeleton compared to the full-width. However, they also showed much greater sensitivity to threshold selection, with very rapid fall-off in performance moving away from the optimal threshold (Fig. 5m). Without a GT for each image, such automatic optimization of the threshold value for any of the algorithms would not be possible, so methods that are highly sensitive to the threshold value are likely to perform much worse when used for typical data sets that lack GTs.

When the above procedure was applied to all leaves in the dataset, the *F*_*β2*_ statistic values for the CNN were higher than for all other algorithms: 0.945 vs 0.837, t=-36.463, df=1062.7, p<10^−16^ (Fig. 6a). Thus, CNN enhancement gave accurate and robust segmentation of these leaf vein networks and out-performed the currently available network enhancement and segmentation methods tested. Performance was also consistent across plant families, and by inference, across diverse network architectures (Fig. 6b).

**Figure 6:**
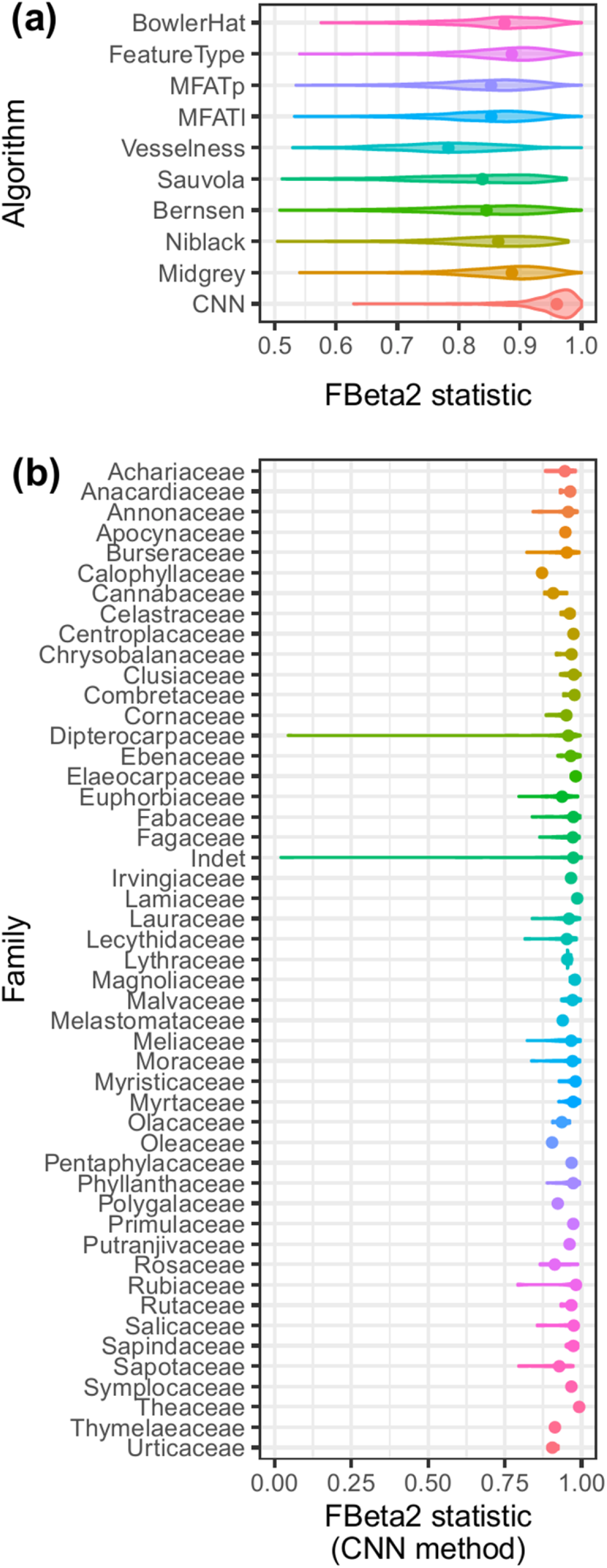
Relative CNN performance for pixel-based PR metrics. (**a**) The *F*_*β2*_ statistic from P-R analysis against FWB-GT tracings was measured using different enhancement algorithms for all 727 specimens using the optimized threshold selection for each method to ensure the best possible performance. (**b**) The ensemble CNN *F*_*β2*_ was also compared across plant families with different network architectures.

As the best PR performance based on pixel classification may not equate to the optimal network extraction, we also examined the impact of varying thresholds on a set of basic network metrics including *Area eccentricity* (Fig. 7a)*, Area mean* (Fig. 7b), *FEV ratio* (Fig. 7c), *No. Junctions* (Fig. 7d), *Length density* (Fig. 7e), and *Node density* (Fig. 7f). For all metrics, the CNN approach showed the lowest fractional error from the GT skeleton, followed by the BowlerHat, Phase-congruency and Hessian-based techniques. The adaptive threshold methods performed very poorly, with fractional errors 1-2 log_e_ units adrift.

**Figure 7:**
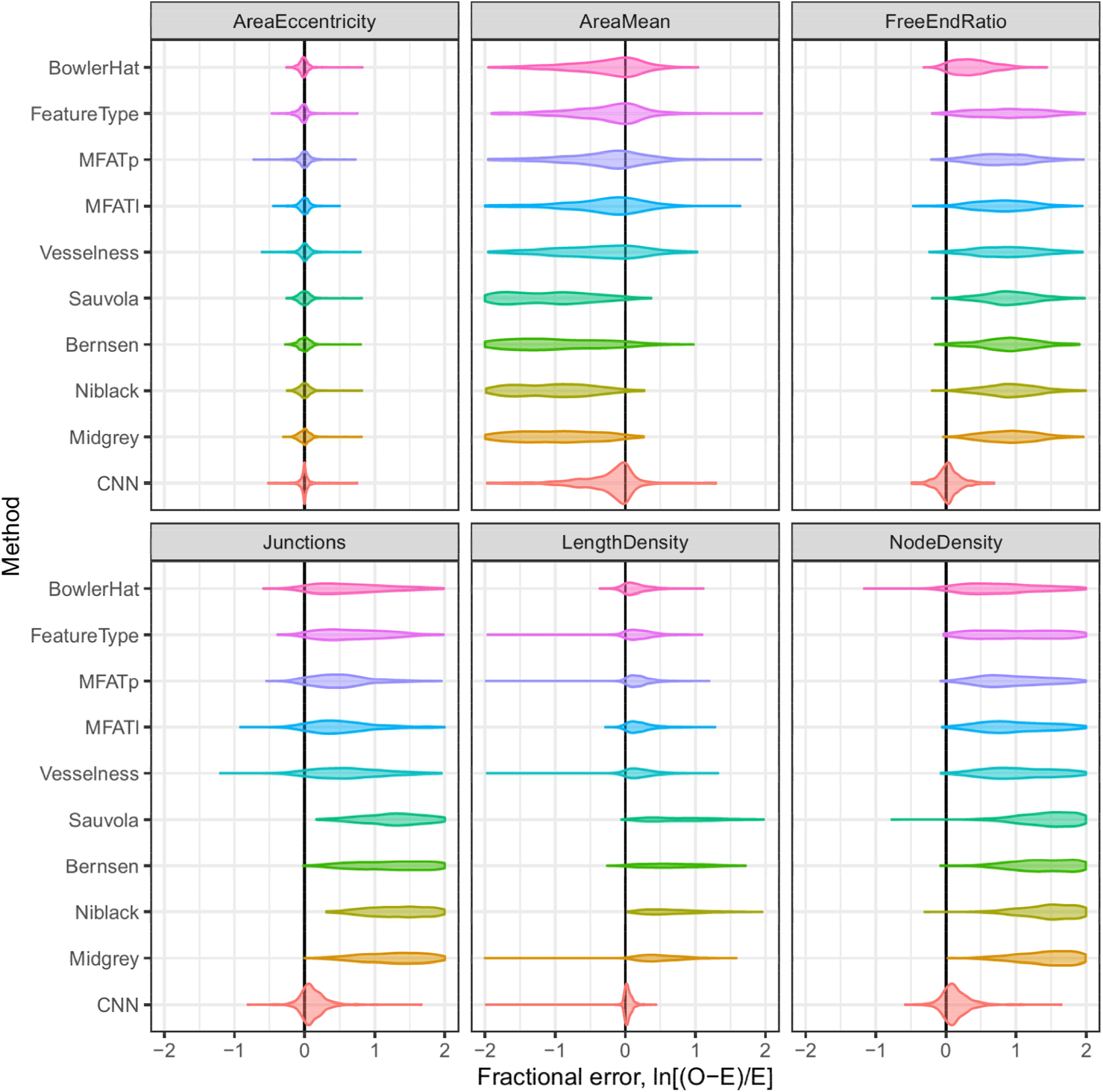
Relative CNN performance for network metrics. The performance of each network enhancement and segmentation method was compared for a set of network metrics, and presented as the log fractional error (log_e_(O-E/E) from the GT skeleton.

### Venation comparisons using single-scale metrics

A range of metrics were calculated following CNN network extraction from the full-size images for the veins, areoles and polygonal regions (Tables S2-5). Each metric had distribution characteristic of the species. For example, the relationship between vein width and length (Fig. 8a) or between loop circularity and elongation (Fig. 8b), showed markedly different patterns between the six species illustrated. For example, in some species (*A. odoratissimus* and *D. lanceolata*), the vein width showed little separation into distinct vein orders, whilst for the others, discrete vein classes were readily identifiable (arrows in Fig. 8a), but with markedly different numbers in each class.

**Figure 8:**
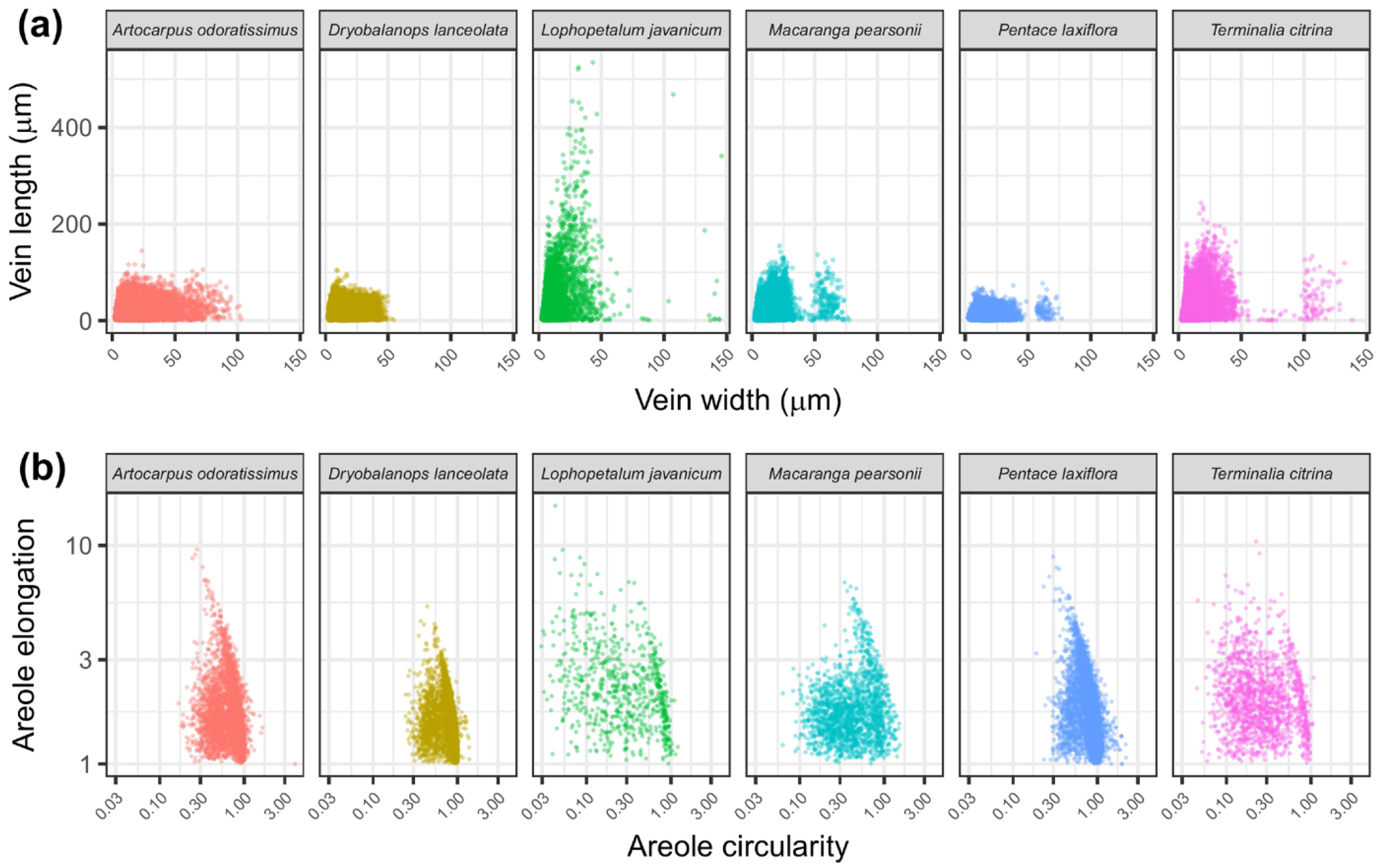
Example single-scale metrics for leaf venation networks. (**a**) Bivariate relationship between vein width and vein length for the six species shown in Fig. 4. Arrows indicate the occurrence of discrete minor vein size classes in some species. (**b**) Bivariate relationship between areole circularity and areole elongation. Points represent each unique vein segment (**a**) or areole (**b**).

### Venation comparisons using HLD and multi-scale metrics

Given the limitations of single scale metrics, we explored analysis using HLD to progressively group regions of the leaf into clusters at different scales to generate multi-scale statistics for each metric (Fig. 3g,h). For example, the spatial scaling of vein density (*VTotLD*) changed as a function of the fusion vein width (*Wid*) (Fig. 9a). At small scales, some species had a much higher vein density (e.g. *P. laxiflora*). However, this reduced rapidly once a specific size class of veins was removed (50μm in this case – Fig. 8a). In some other species (e.g. *T. citrina*), as the scale increased (or equivalently as the size class of the intervening vein increased), the vein density was maintained to a much greater extent.

**Figure 9:**
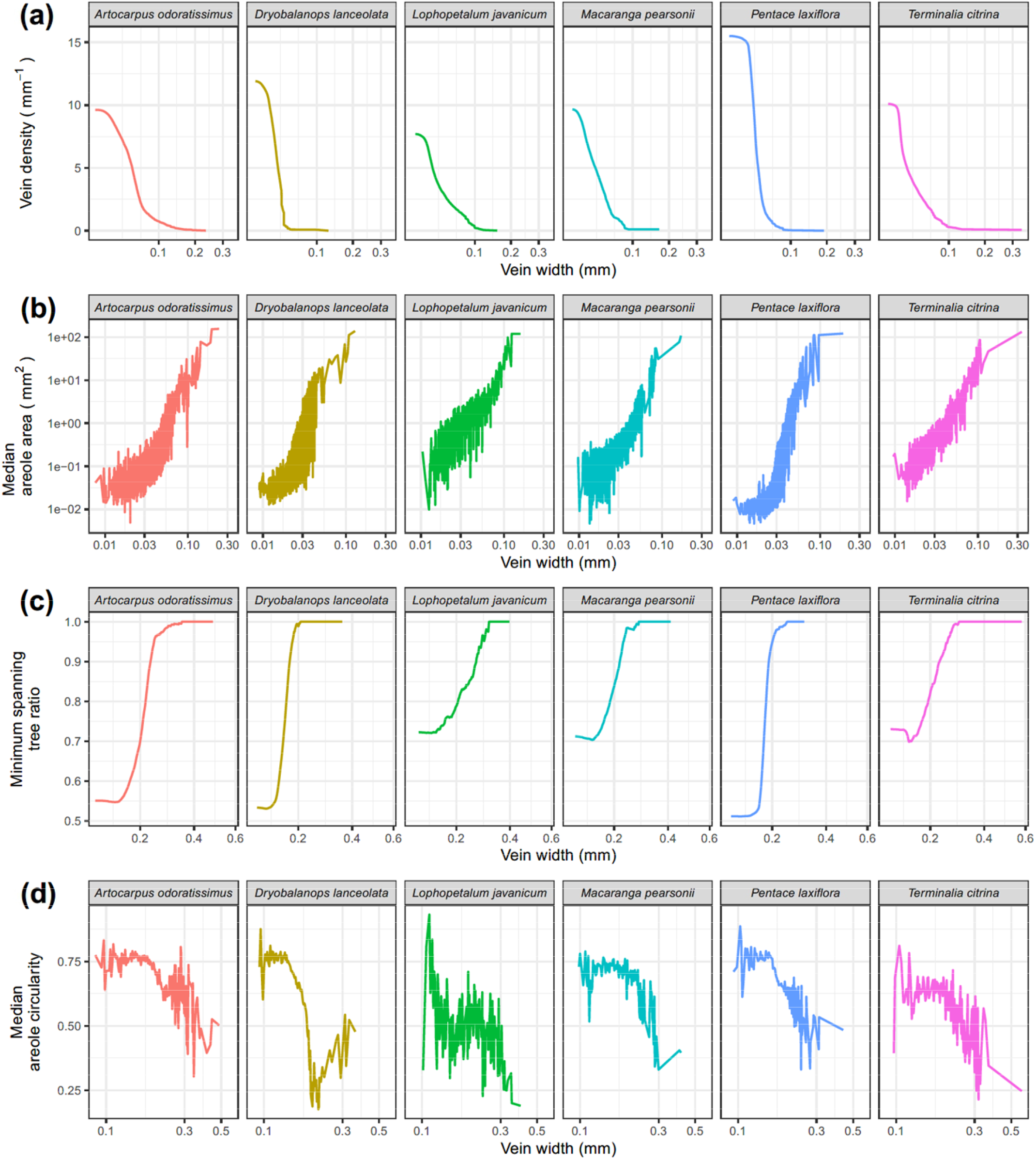
Example multi-scale metrics for leaf venation networks. (**a**) Changes in vein density with increasing vein width for the six species shown in Figure 4 following successive vein removal during HLD. (**b**) Changes in median areole area with increasing vein width removal. (**c**) Changes in minimum spanning tree ratio with increasing vein width removal. (**d**) Changes in median areole circularity with increasing areole area following fusion.

The different vein sizes and overall vein density reflected different partitioning of the leaf lamina (Fig. 9b). Thus, the median loop area was an order of magnitude larger for *L. javanicum, M. pearsonii* and *T. citrina*, relative to other species at small scales, indicating a lower areole density in these species. In contrast, at intermediate scales, other species like *D. lanceolata* or *P. laxiflora* transitioned rapidly to low levels of looping from a dense initial value, whilst *A. odoratissimus* maintained a more consistent response across scales, reflecting the relative investment in loops and redundant flow pathways at different spatial scales.

The level of redundancy was also reflected in the amount of network branching across spatial scales measured by the minimum spanning tree (MST) ratio (Fig. 9c). The MST ratio increased with vein size class fastest for *D. lanceolata* and *P. laxiflora*, and slowest for *L. javanicum*. In all cases there were shifts in gradient corresponding to scale transitions between veins. *T. citrina* and *M. pearsonii* also showed non-monotonic changes in the MST ratio, consistent with early pruning of FEV and tree-like branches to give a minimum at an intermediate spatial scale. Thus, this multi-scale statistic indicates how investment in veins of different sizes yields different branching organisation.

The space-filling geometry of the network also changed across spatial scales (Fig. 9d). Thus, the median areole circularity decreased with vein size class in all species, but at different rates. Areole circularity was similar between species at small spatial scales, but at larger spatial scales. *D. lanceolata* had the least circular areoles due to fusing regions becoming more elongated with a more parallel vein architecture, while *A. odoratissimus* maintained the most circular areoles.

## Discussion

### CNNs improve network segmentation accuracy and robustness

Ensemble CNNs provide an alternative approach for leaf vein network segmentation with high accuracy, as judged both by P-R analysis against GT tracings using pixel classification and a set of network measures. Ensemble CNNs also provided consistently better segmentation of leaf vein networks from field-collected leaves across many different species and leaf vein architectures than other currently available network extraction algorithms. In addition, the CNN probability map produced smooth, full-width segmentation of veins, even in the presence of image artifacts such as differential clearing, air bubbles, or trichomes. This in turn led to improved robustness, precision and accuracy in graph extraction, vein width estimation, and subsequent quantitative metrics.

We note that this deep learning approach contrasts with previous neural network approaches for leaf architecture (Grinblat *et al.*, 2016), which used simpler network architectures and more limited training data, with a goal to develop a classification system rather than extract the venation network itself.

Ensemble CNNs allowed rapid characterization of minor vein networks at multiple spatial scales with minimal manual intervention, removing a major bottleneck of manual tracing. For comparison, hand-tracing the equivalent area predicted by the CNNs would have required >36,000 person-hours to complete. Nevertheless, the results presented here are truncated by the leaf sample extents, which prevented exploration of larger scales than ~10 × 10 mm^2^ areas. The trained ensemble CNNs used input images with pixel resolution (1.68 μm pix^−1^) and contrast adjusted to cover the full dynamic range following CLAHE, and can be used without modification to analyse other leaf images of comparable contrast and resolution. Other leaf vein images at lower spatial resolution can still be segmented well using up-sampling with interpolation, to ensure that the smallest veins are ~5 pixels wide. A typical example is shown in Fig. S9 for a leaf originally collected at 6.7 μm pix^−1^ using the LeafVeinCNN GUI (Fig. S2). Nevertheless, the performance of the ensemble CNNs is driven by the quality of the GT and the input images. For example, the LeafVeinCNN may have limited performance on lower-resolution images that are typical for digitized historic cleared leaf collections, e.g. the Smithsonian / Wolfe collection (Lobet *et al.*, 2013; Das *et al.*, 2014), without additional training, or using approaches such as generative adversarial networks (GANs) to refine segmentations and avoid unlikely predictions that have large blotches or disconnected areas.

### Network analysis across multiple scales

Most studies have primarily focused on descriptors of the network at single scales, e.g. the density of major and minor veins (Sack *et al.*, 2012), or metrics for the shape of areoles (Blonder *et al.*, 2011; Price & Weitz, 2014). At multiple scales, other studies have suggested that the hierarchical nesting of veins and areoles may distinguish species or have functional significance (Katifori *et al.*, 2010; Katifori & Magnasco, 2012; Ronellenfitsch *et al.*, 2015; Brodribb *et al.*, 2016; Fiorin *et al.*, 2016). Existing systematic manuals recognize a broad diversity of leaf venation forms and associated terminology that are not well-captured by extant quantitative metrics (Ellis *et al.*, 2009). Additionally, prior efforts to leverage HLD yielded a set of statistics, such as Strahler number, derived originally from analysis of river basins, which are quite difficult to interpret in terms of biological processes (Pelletier & Turcotte, 2000; Price *et al.*, 2011; Ronellenfitsch *et al.*, 2015; Lasser & Katifori, 2017). Here, we also used HLD to determine scaling relationships, but retained a set of more conventional metrics to compare network architecture across scales. These multi-scale statistics may provide additional insights into how network architecture influences transport, support, defense, embolism resistance, and resilience to herbivory (Sack *et al.*, 2008; Katifori *et al.*, 2010; Brodribb *et al.*, 2016; Blonder *et al.*, 2018). Indeed, we recently have made such an analysis of these network–function linkages using networks extracted from this dataset (Blonder, 2020), and we propose that multi-scale statistics may enable a more quantitative and richer description of network architecture that supplements the utility of qualitative classifications.

## Acknowledgments

We thank David Ford for assistance with the GPU cluster. This work was supported by the Human Frontier Science Program (RGP0053/2012, MDF), the Leverhulme Trust (RPG-2015-437, MDF), the UK Natural Environment Research Council (NERC) (NE/M019160/1, BB), the US National Science Foundation (DEB-2025282, BB), and an ERC Advanced Investigator Award (GEM-TRAIT 321131, YM). The field work was funded by the UK NERC BALI consortium (NE/K016253/1). The SAFE Project was funded by the Sime Darby Foundation and the UK NERC.

## Author contributions

HX developed the CNN training algorithms. MDF developed the network extraction and analysis algorithms, and GUI interface. MJ chemically cleared and imaged the leaves. BB carried out the fieldwork and the statistical analysis. YM and BB secured funding for the field work and lab work, and computing infrastructure. HX, MDF, and BB contributed equally to the manuscript.

## Supporting Online Material

**Supplementary Figure 1: Schematic illustration of the network metrics extracted.**

**Supplementary Figure 2: The LeafVeinCNN graphical user interface (GUI)**

**Supplementary Figure 3: Sensitivity analysis for varying levels of contrast, with no correction.**

**Supplementary Figure 4: Sensitivity analysis for varying levels of contrast followed by CLAHE**

**Supplementary Figure 5: Sensitivity Analysis for varying levels of Contrast-Limited Adaptive Histogram Equalization (CLAHE)**

**Supplementary Figure 6: Effect of increasing blur on ensemble CNN performance**

**Supplementary Figure 7: Effect of image resolution on ensemble CNN performance**

**Supplementary Figure 8: Effect of up-sampling to correct for low resolution input images**

**Supplementary Figure 9: Application of LeafVeinCNN to an intact leaf**.

**Supplementary Table 1: Image enhancement processing parameters**

**Supplementary Table 2: Vein metrics**

**Supplementary Table 3: Node metrics**

**Supplementary Table 4: Areole metrics**

**Supplementary Table 5: Polygonal region metrics**

**Supplementary Table 6: HLD metrics.**

**Supplementary Table 7: Summary statistics for vein, node, areole and polygon metrics**

## References

Alhasson HF, Alharbi SS, Obara B 2018. 2D and 3D Vascular Structures Enhancement via Multiscale Fractional Anisotropy Tensor. Proceedings of the European Conference on Computer Vision (ECCV). 0–0.

Berens P. 2009. CircStat: a MATLAB toolbox for circular statistics. J. Stat. Softw. 31: 1–21.

Bernsen J 1986. Dynamic thresholding of gray-level images. Proc. ICAR. Berlin. 1251–1255.

Blonder B, Both S, Jodra M, Majalap N, Burslem D, Teh YA, Malhi Y. 2019. Leaf venation networks of Bornean trees: images and hand-traced segmentations. Ecology 100: e02844.

Blonder B, Both, S., Jodra, M., Xu, H., Fricker, M.D., Matos, I., Majalap-Lee, N., Burslem, D., T, Y. A., Malhi, Y. 2020. Linking functional traits to multiscale statistics of leaf venation networks. New Phytol. in press.

Blonder B, Carlo F, Moore J, Rivers M, Enquist BJ. 2012. X-ray imaging of leaf venation networks. New Phytol. 196: 1274–1282.

Blonder B, Enquist BJ. 2014. Inferring climate from angiosperm leaf venation networks. New Phytol. 204: 116–126.

Blonder B, Royer DL, Johnson KR, Miller I, Enquist BJ. 2014. Plant Ecological Strategies Shift Across the Cretaceous–Paleogene Boundary. PLOS Biology 12: e1001949.

Blonder B, Salinas N, Patrick Bentley L, Shenkin A, Chambi Porroa PO, Valdez Tejeira Y, Boza Espinoza TE, Goldsmith GR, Enrico L, Martin R. 2018. Structural and defensive roles of angiosperm leaf venation network reticulation across an Andes-Amazon elevation gradient. J. Ecol. 106: 1683–1699.

Blonder B, Violle C, Bentley LP, Enquist BJ. 2011. Venation networks and the origin of the leaf economics spectrum. Ecol. Lett. 14: 91–100.

Bohn S, Andreotti B, Douady S, Munzinger J, Couder Y. 2002. Constitutive property of the local organization of leaf venation networks. Phys. Rev. E 65: 061914.

Both S, Riutta T, Paine CT, Elias DM, Cruz R, Jain A, Johnson D, Kritzler UH, Kuntz M, Majalap-Lee N. 2018. Logging and soil nutrients independently explain plant trait expression in tropical forests. New Phytol.

Boyce CK, Brodribb T, Feild TS, Zwieniecki MA. 2009. Angiosperm leaf vein evolution was physiologically and environmentally transformative. Proc. Roy. Soc. Lond. B 276: 1771–1776.

Bresenham JE. 1965. Algorithm for computer control of a digital plotter. IBM Systems Journal 4: 25–30.

Brodribb T, Feild T, Jordan G. 2007. Leaf maximum photosynthetic rate and venation are linked by hydraulics. Plant Physiol. 144: 1890.

Brodribb T, Feild TS. 2010. Leaf hydraulic evolution led a surge in leaf photosynthetic capacity during early angiosperm diversification. Ecol. Lett. 13: 175–183.

Brodribb TJ, Bienaimé D, Marmottant P. 2016. Revealing catastrophic failure of leaf networks under stress. Proc. Natl. Acad. Sci. USA 113: 4865–4869.

Brodribb TJ, Feild TS, Sack L. 2010. Viewing leaf structure and evolution from a hydraulic perspective. Functional Plant Biology 37: 488–498.

Bühler J, Rishmawi L, Pflugfelder D, Huber G, Scharr H, Hülskamp M, Koornneef M, Schurr U, Jahnke S. 2015. phenoVein—A tool for leaf vein segmentation and analysis. Plant Physiol. 169: 2359–2370.

Burt P, Adelson E. 1983. The Laplacian Pyramid as a Compact Image Code. IEEE Trans. Comm. 31: 532–540.

Carvalho MR, Losada JM, Niklas KJ. 2018. Phloem networks in leaves. Curr. Opin. Plant Biol. 43: 29–35.

Carvalho MR, Turgeon R, Owens T, Niklas KJ. 2017. The hydraulic architecture of Ginkgo leaves. Am. J. Bot. 104: 1285–1298.

Das A, Bucksch A, Price CA, Weitz JS. 2014. ClearedLeavesDB: an online database of cleared plant leaf images. Plant methods 10: 8–8.

de Boer HJ, Eppinga MB, Wassen MJ, Dekker SC. 2012. A critical transition in leaf evolution facilitated the Cretaceous angiosperm revolution. Nat. Comm. 3: 1221.

Dhondt S, Van Haerenborgh D, Van Cauwenbergh C, Merks RMH, Philips W, Beemster GTS, Inzé D. 2012. Quantitative analysis of venation patterns of Arabidopsis leaves by supervised image analysis. Plant J. 69: 553–563.

Dirnberger M, Kehl T, Neumann A. 2015. NEFI: Network Extraction From Images. Scientific Reports 5: 15669.

Dodds PS. 2010. Optimal form of branching supply and collection networks. Phys. Rev. Lett. 104: 048702.

Ellis B, Daly DC, Hickey LJ, Johnson KR, Mitchell JD, Wilf P, Wing SL. 2009. Manual of Leaf Architecture: Cornell University Press.

Fiorin L, Brodribb TJ, Anfodillo T. 2016. Transport efficiency through uniformity: organization of veins and stomata in angiosperm leaves. New Phytol. 209: 216–227.

Frangi A, Niessen W, Vincken K, Viergever M 1998. Multiscale vessel enhancement filtering. In: Wells W, Colchester A, Delp S eds. Medical Image Computing and Computer-Assisted Interventation: Springer Berlin / Heidelberg, 130–137.

Fricker MD, Akita D, Heaton LLM, Jones N, Obara B, Nakagaki T. 2017. Automated analysis of *Physarum* network structure and dynamics. J. Physics D 50: 254005.

Gan Y, Rong Y, Huang F, Hu L, Yu X, Duan P, Xiong S, Liu H, Peng J, Yuan X. 2019. Automatic hierarchy classification in venation networks using directional morphological filtering for hierarchical structure traits extraction. Computational Biology and Chemistry 80: 187–194.

Glorot X, Bengio Y 2010. Understanding the difficulty of training deep feedforward neural networks.In Yee Whye T, Mike T. Proceedings of the Thirteenth International Conference on Artificial Intelligence and Statistics. Proceedings of Machine Learning Research: PMLR. 249--256.

Glorot X, Bordes A, Bengio Y 2011. Deep sparse rectifier neural networks. Proc. AISTATS. 315–323.

Green WA, Little SA, Price CA, Wing SL, Smith SY, Kotrc B, Doria G. 2014. *Reading the leaves: A comparison of leaf rank and automated areole measurement* for quantifying aspects of leaf venation. Applications in Plant Sciences 2: apps.1400006.

Grinblat GL, Uzal LC, Larese MG, Granitto PM. 2016. Deep learning for plant identification using vein morphological patterns. Comput. Electron. Agric. 127: 418–424.

Hansen LK, Salamon P. 1990. Neural network ensembles. IEEE Trans. Pattern Anal. Mach. Intell. 12: 993–1001.

Hickey LJ 1979. A revised classification of the architecture of dicotyledonous leaves. In: Metcalfe CR, Chalk L eds. Anatomy of the Dicotyledons. Vol 1, Systematic Anatomy of the Leaf and Stem. Oxford: Clarendon Press, 25–39.

Ioffe S, Szegedy C. 2015. Batch Normalization: Accelerating Deep Network Training by Reducing Internal Covariate Shift. CoRR arXiv:1502.03167v3.

John GP, Scoffoni C, Buckley TN, Villar R, Poorter H, Sack L. 2017. The anatomical and compositional basis of leaf mass per area. Ecol. Lett. 20: 412–425.

Kang J, Dengler N. 2004. Vein pattern development in adult leaves of Arabidopsis thaliana. Int. J. Plant Sci. 165: 231–242.

Katifori E. 2018. The transport network of a leaf. Comptes Rendus Physique 19: 244–252.

Katifori E, Magnasco MO. 2012. Quantifying loopy network architectures. PloS One 7: e37994.

Katifori E, Szollosi GJ, Magnasco MO. 2010. Damage and fluctuations induce loops in optimal transport networks. Phys. Rev. Lett 104: 048704.

Kovesi PD. 1999. Image features from phase congruency. Videre 1: 1–26.

Kovesi PD. 2000. MATLAB and Octave functions for computer vision and image processing. http://www.csse.uwa.edu.au/~pk/research/matlabfns/.

Krizhevsky A, Sutskever I, Hinton GE 2012. Imagenet classification with deep convolutional neural networks. Adv. Neural Inf. Process Syst. 1097–1105.

Larese MG, Namías R, Craviotto RM, Arango MR, Gallo C, Granitto PM. 2014. Automatic classification of legumes using leaf vein image features. Pattern Recognition 47: 158–168.

Lasser J, Katifori E. 2017. NET: a new framework for the vectorization and examination of network data. Source Code for Biology and Medicine 12: 4.

LeCun Y, Bengio Y, Hinton G. 2015. Deep learning. Nature 521: 436–444.

Lobet G, Draye X, Périlleux C. 2013. An online database for plant image analysis software tools. Plant Methods 9: 1–8.

Long J, Shelhamer E, Darrell T 2015. Fully convolutional networks for semantic segmentation. Proceedings of the IEEE conference on computer vision and pattern recognition. 3431–3440.

Lopez-Molina C, De Baets B, Bustince H. 2013. Quantitative error measures for edge detection. Pattern Recognition 46: 1125–1139.

Manze U. 1967. Die Nervaturdichte der Blätter als Hilfsmittel der Paläoklimatologie. Geologisches Institut der Universität zu Köln.

McKown AD, Cochard H, Sack L. 2010. Decoding leaf hydraulics with a spatially explicit model: principles of venation architecture and implications for its evolution. American Naturalist 175: 447–460.

Mileyko Y, Edelsbrunner H, Price CA, Weitz JS. 2012. Hierarchical Ordering of Reticular Networks. PLoS ONE 7: e36715.

Mounsef J, Karam L 2012. Fully automated quantification of leaf venation structure. Proc. ICAI: WorldComp. 1.

Niblack W. 1985. An Introduction to Digital Image Processing: Strandberg Publishing Company.

Niklas KJ. 1999. A mechanical perspective on foliage leaf form and function. New Phytol. 143: 19–31.

Obara B, Grau V, Fricker MD. 2012. A bioimage informatics approach to automatically extract complex fungal networks. Bioinformatics 28: 2374–2381.

Pain C, Kriechbaumer V, Kittelmann M, Hawes C, Fricker M. 2019. Quantitative analysis of plant ER architecture and dynamics. Nat. Comm. 10: 984.

Parsons-Wingerter P, Vickerman MB, Paul A-L, Ferl RJ. 2014. Mapping by VESGEN of leaf venation patterning in Arabidopsis with bioinformatic dimensions of gene expression. Gravitational and Space Research 2.

Pelletier JD, Turcotte DL. 2000. Shapes of river networks and leaves: are they statistically similar? Philosophical Transactions of the Royal Society of London. Series B: Biological Sciences 355: 307–311.

Pérez-Harguindeguy N, Díaz S, Garnier E, Lavorel S, Poorter H, Jaureguiberry P, Bret-Harte M, Cornwell W, Craine J, Gurvich D. 2013. New handbook for standardised measurement of plant functional traits worldwide. Aust. J. Bot. 61: 167–234.

Price CA, Knox S-JC, Brodribb TJ. 2014. The Influence of Branch Order on Optimal Leaf Vein Geometries: Murray’s Law and Area Preserving Branching. PLOS ONE 8: e85420.

Price CA, Symonova O, Mileyko Y, Hilley T, Weitz JS. 2011. Leaf Extraction and Analysis Framework Graphical User Interface: Segmenting and Analyzing the Structure of Leaf Veins and Areoles. Plant Physiol. 155: 236–245.

Price CA, Weitz JS. 2014. Costs and benefits of reticulate leaf venation. BMC Plant Biol. 14: 234.

Price CA, Wing S, Weitz JS. 2012. Scaling and structure of dicotyledonous leaf venation networks. Ecol. Lett. 15: 87–95.

Riutta T, Malhi Y, Kho LK, Marthews TR, Huaraca Huasco W, Khoo M, Tan S, Turner E, Reynolds G, Both S. 2018. Logging disturbance shifts net primary productivity and its allocation in Bornean tropical forests. Glob. Change Biol. 24: 2913–2928.

Rolland-Lagan A-G, Amin M, Pakulska M. 2009. Quantifying leaf venation patterns: two-dimensional maps. Plant J. 57: 195–205.

Ronellenfitsch H, Lasser J, Daly DC, Katifori E. 2015. Topological phenotypes constitute a new dimension in the phenotypic space of leaf venation networks. PLoS Computational Biology 11: e1004680.

Ronneberger O, Fischer P, Brox T 2015. U-Net: Convolutional Networks for Biomedical Image Segmentation: Springer International Publishing. 234–241.

Roth-Nebelsick A, Uhl D, Mosbrugger V, Kerp H. 2001. Evolution and function of leaf venation architecture: a review. Ann. Bot. 87: 553–566.

Rother C, Kolmogorov V, Blake A 2004. Grabcut: Interactive foreground extraction using iterated graph cuts. ACM transactions on graphics (TOG): ACM. 309–314.

Sack L, Dietrich EM, Streeter CM, Sánchez-Gómez D, Holbrook NM. 2008. Leaf palmate venation and vascular redundancy confer tolerance of hydraulic disruption. Proceedings of the National Academy of Sciences 105: 1567–1572.

Sack L, Frole K. 2006. Leaf structural diversity is related to hydraulic capacity in tropical rain forest trees. Ecology 87: 483–491.

Sack L, Scoffoni C. 2013. Leaf venation: structure, function, development, evolution, ecology and applications in the past, present and future. New Phytol. 198: 983–1000.

Sack L, Scoffoni C, McKown AD, Frole K, Rawls M, Havran JC, Tran H, Tran T. 2012. Developmentally based scaling of leaf venation architecture explains global ecological patterns. Nat. Comm. 3: 837.

Saito T, Rehmsmeier M. 2015. The Precision-Recall plot is more informative than the ROC plot when evaluating binary classifiers on imbalanced datasets. PLOS ONE 10: e0118432.

Sauvola J, Pietikäinen M. 2000. Adaptive document image binarization. Pattern Recognition 33: 225–236.

Sazak Ç, Nelson CJ, Obara B. 2018. The multiscale bowler-hat transform for blood vessel enhancement in retinal images. Pattern Recognition.

Schneider JV, Rabenstein R, Wesenberg J, Wesche K, Zizka G, Habersetzer J. 2018. Improved non-destructive 2D and 3D X-ray imaging of leaf venation. Plant methods 14: 7.

Sharon E, Sahaf M 2018. The Mechanics of Leaf Growth on Large Scales. In: Geitmann A, Gril J eds. Plant Biomechanics: From Structure to Function at Multiple Scales. Cham: Springer International Publishing, 109–126.

Simonyan K, Zisserman A. 2014. Very deep convolutional networks for large-scale image recognition. arXiv:1409.1556v6.

Sollich P, Krogh A. 1996. Learning with ensembles: How overfitting can be useful. Adv. Neural Inf. Process Syst. 8: 190–196.

Trivett ML, Pigg KB 1996. A survey of reticulate venation among fossil and living land plants. Flowering Plant Origin, Evolution & Phylogeny: Springer, 8–31.

Uhl D, Mosbrugger V. 1999. Leaf venation density as a climate and environmental proxy: a critical review and new data. Palaeogeography, Palaeoclimatology, Palaeoecology 149: 15–26.

Werbos P. 1974. Beyond Regression: New Tools for Prediction and Analysis in the Behavioral Sciences. Harvard University.

Wing SL. 1992. High-Resolution Leaf X-Radiography in Systematics and Paleobotany. Am. J. Bot. 79: 1320–1324.

Zhang TY, Suen CY. 1984. A fast parallel algorithm for thinning digital patterns. Comm. ACM 27: 236–239.

Zuiderveld K. 1994. Contrast limited adaptive histogram equalization. Graphics gems: 474–485.

