## Supplementary Information for "Automated and accurate segmentation of leaf venation networks via deep learning"

New Phytologist

***New Phytologist* Supporting Information**

Article title: Automated and accurate segmentation of leaf venation networks via deep learning Supplementary Information
Authors: H. Xu, B. Blonder, M. Jodra, Y. Malhi, and M.D. Fricker.
Article acceptance date: Click here to enter a date.

The following Supporting Information is available for this article:

**Fig. S1** Schematic illustration of the network metrics extracted.

**Fig. S2** The LeafVeinCNN graphical user interface (GUI).

**Fig. S3** Sensitivity analysis for varying levels of contrast, with no correction.

**Fig. S4** Sensitivity analysis for varying levels of contrast followed by CLAHE.

**Fig. S5** Sensitivity Analysis for varying levels of Contrast-Limited Adaptive Histogram

Equalization (CLAHE)

**Fig. S6** Effect of increasing blur on ensemble CNN performance

**Fig. S7** Effect of image resolution on ensemble CNN performance

**Fig. S8** Effect of up-sampling to correct for low resolution input images.

**Fig. S9** Application of LeafVeinCNN to an intact leaf.

**Table S1** Image enhancement processing parameters

**Table S2** Vein metrics

**Table S3** Node metrics

**Table S4** Areoles metrics.

**Table S5** Polygonal region metrics

**Table S6** HLD metrics

**Table S7** Summary statistics for vein, node, areole and polygon metrics

**Fig. S1 Schematic illustration of the network metrics extracted**. (**a**) A stylised part of the leaf vein network containing a fully enclosed areole. (**b**) The extracted pixel skeleton showing loops (magenta) and tree-like (green) components of the skeleton. (**c**) The network following conversion to a weighted graph with nodes connected by edges. (**d**) Representation of individual vein metrics. The distribution for the complete sample can be summarized by different statistics including mean, median, SD or skewness. (**e**) Node metrics for a typical three-way branch. (**f**) Metrics for areoles (lamina excluding the veins) and polygonal areas (lamina separated by the pixel skeleton). Note all metrics illustrated in (**f**) are calculated for both areoles and polygonal regions in single scale analysis. During HLD, only metrics for the polygonal regions are calculated.

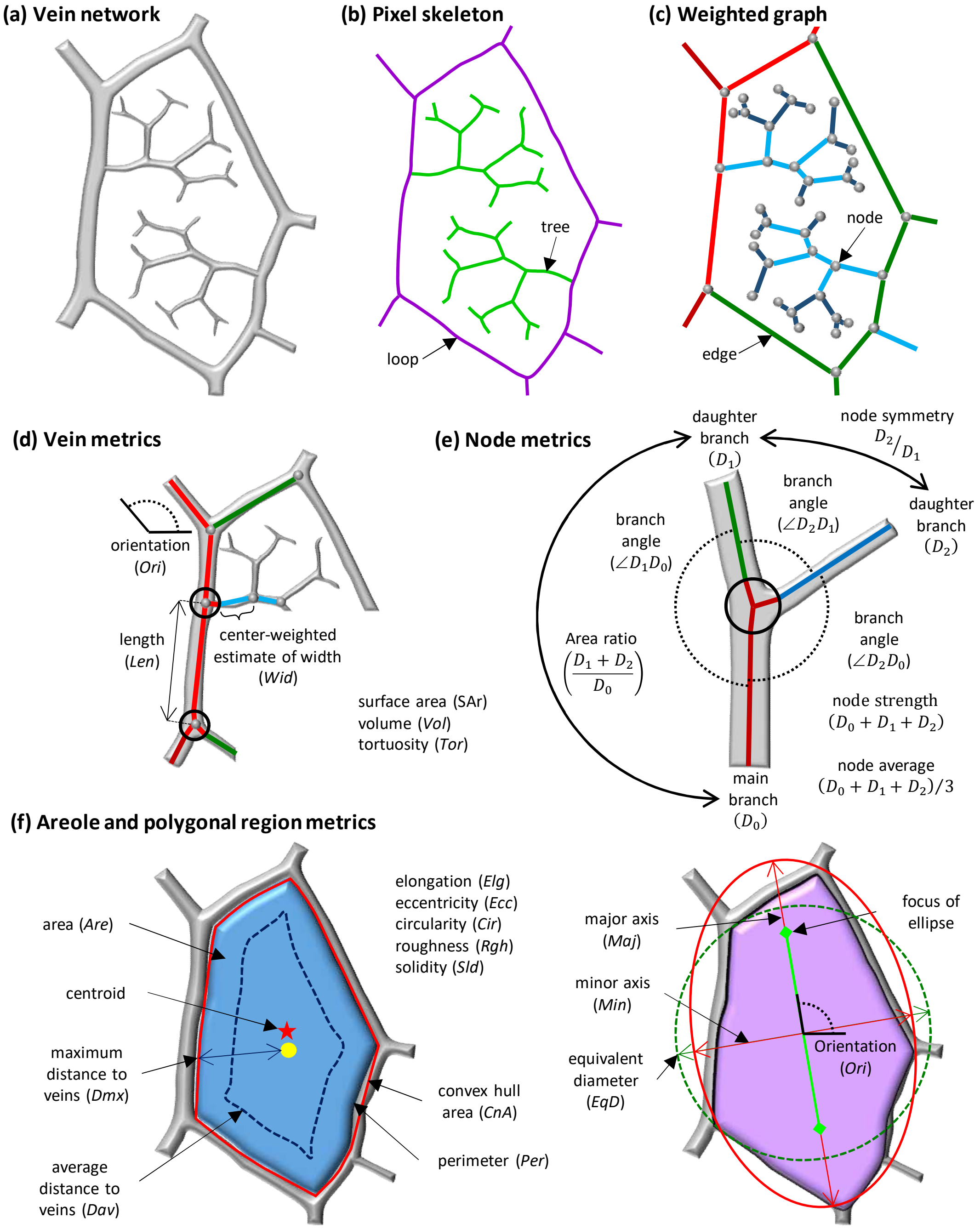

**Fig. S2 The LeafVeinCNN graphical user interface (GUI) and manual.** The LeafVeinCNN GUI runs as a standalone program or within the MATLAB environment and can be used to extract the vein network from any cleared leaf image using the pre-trained CNNs. The manual can be assessed at (URL available on acceptance)

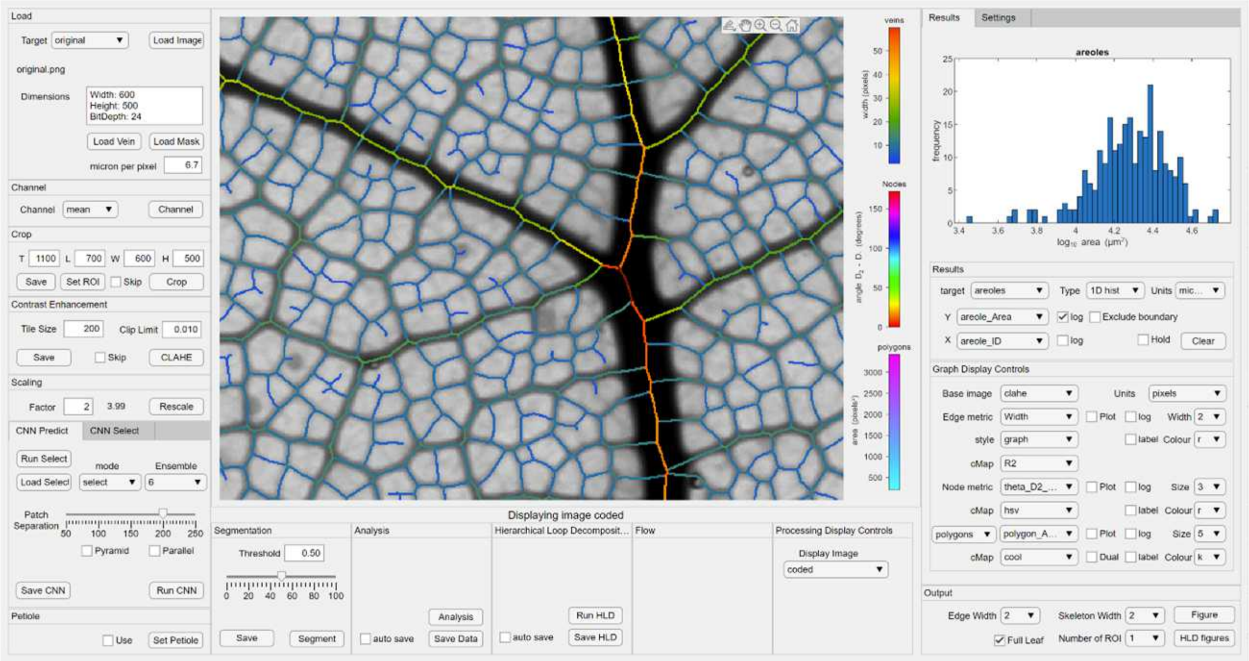

**Fig. S3 : Sensitivity analysis for varying levels of contrast, with no correction.** A small (~3mm x 2mm) section of a leaf image was processed to reduce the contrast range (**a**). The resultant images were enhanced using an ensemble of three CNNs to give a vein prediction (**b**). The mean CNN predictions were thresholded to give a full-width binary (FWB) image and evaluated against a manually-defined FWB-GT image (**c**), or the resultant single-pixel GT-skeleton (**d**). In each case, true positives ($T_P$) are coded in green, false negatives ($F_N$) in red, and false positives ($F_P$) in blue. Precision-Recall (P-R) plots illustrate the performance of each method as the segmentation threshold is varied for the FWB comparison (**e**) and following skeletonisation (**f**). The value of the Dice Similarity Coefficient ($F_1$, asterisk), and $F_{\beta 2}$ statistic (open circle) are shown on the P-R plots and above the corresponding image. The performance of the CNN deteriorates quite badly as the contrast is reduced.

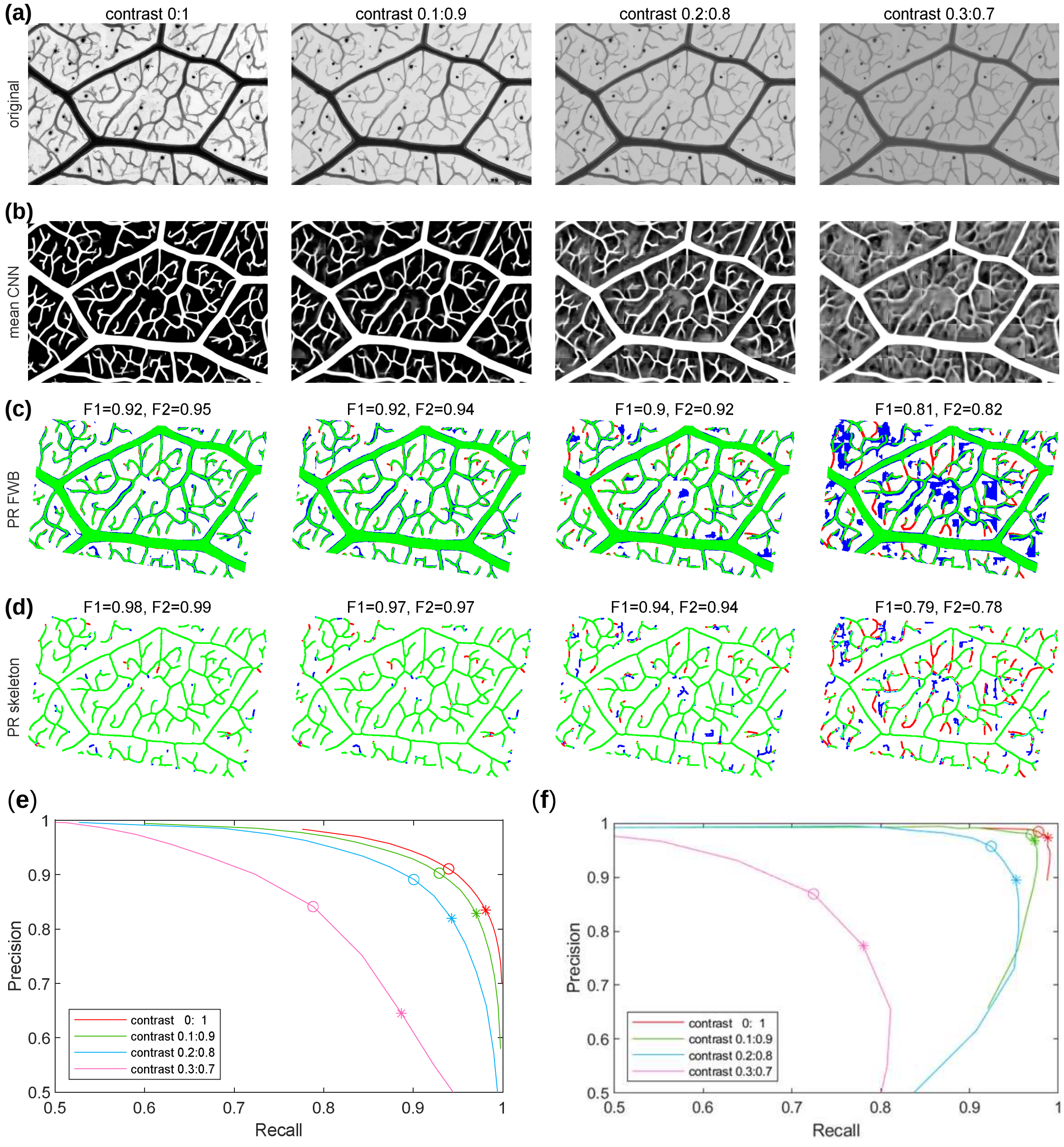

**Fig. S4 Sensitivity analysis for varying levels of contrast followed by CLAHE.** A small (~3mm x 2mm) section of a leaf image was processed to reduce the contrast range, followed by CLAHE with a fixed clip-limit of 0.01 to restore the contrast (**a**). The resultant images were enhanced using an ensemble of three CNNs to give a vein prediction (**b**). The mean CNN predictions were thresholded to give a full-width binary (FWB) image and evaluated against a manually-defined FWB-GT image (**c**), or the resultant single-pixel GT-skeleton (**d**). In each case, true positives ($T_P$) are coded in green, false negatives ($F_N$) in red, and false positives ($F_P$) in blue. Precision-Recall (P-R) plots illustrate the performance of each method as the segmentation threshold is varied for the FWB comparison (**e**) and following skeletonisation (**f**). The value of the Dice Similarity Coefficient ($F_1$, asterisk), and $F_{\beta 2}$ statistic (open circle) are shown on the P-R plots and above the corresponding image. The response of the CNN to low contrast initially can be recovered completely by CLAHE.

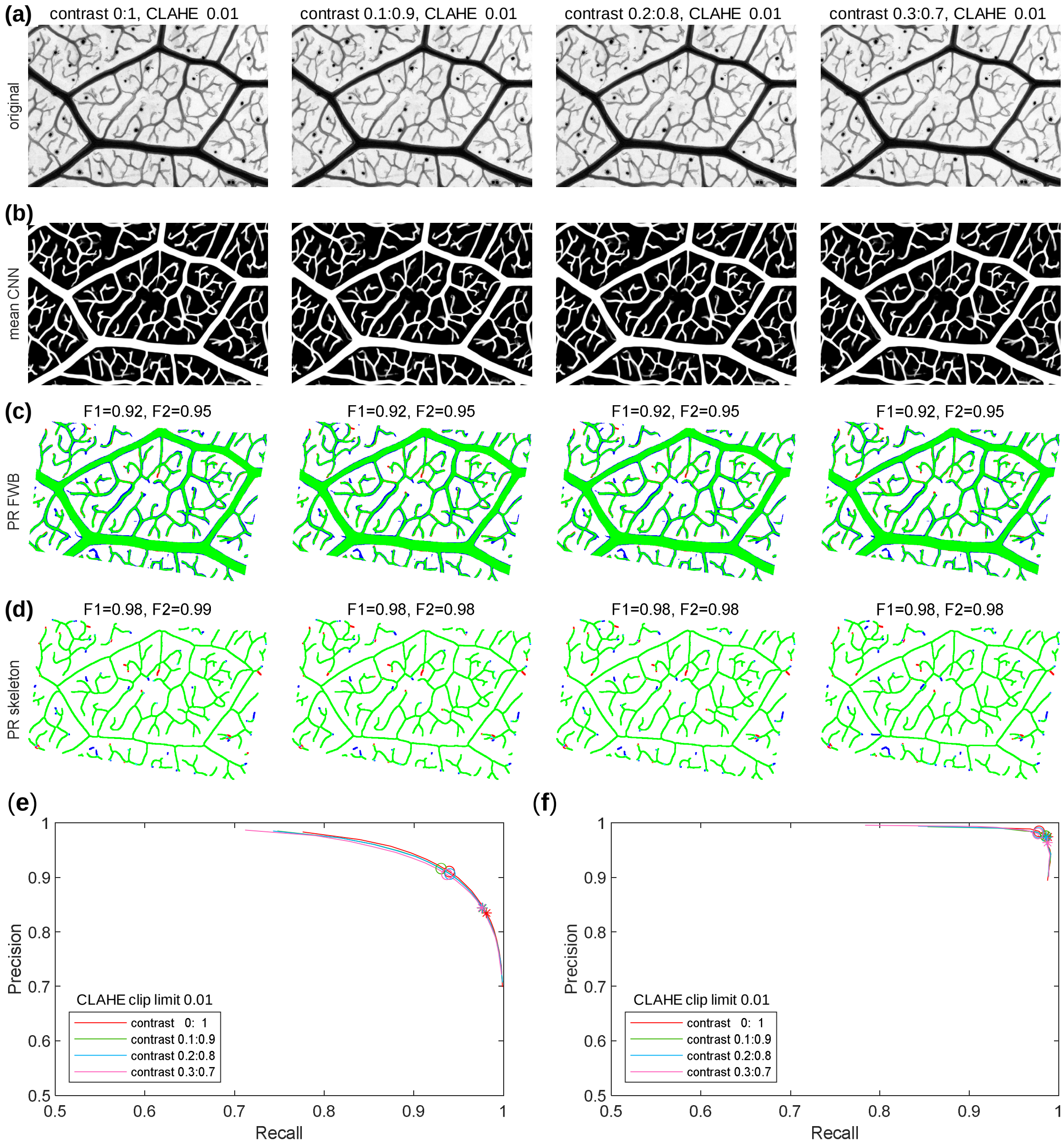

**Fig. S5 Sensitivity Analysis for varying levels of Contrast-Limited Adaptive Histogram Equalization (CLAHE).** A small (~3mm x 2mm) section of a leaf image was processed using CLAHE with varying clip-limit to control the overall contrast (**a**). The resultant images were enhanced using an ensemble of three CNNs to give a vein prediction (**b**). The mean CNN predictions were thresholded to give a full-width binary (FWB) image and evaluated against a manually-defined FWB-GT image (**c**), or the resultant single-pixel GT-skeleton (**d**). In each case, true positives ($T_P$) are coded in green, false negatives ($F_N$) in red, and false positives ($F_P$) in blue. Precision-Recall (P-R) plots illustrate the performance of each method as the segmentation threshold is varied for the FWB comparison (**e**) and following skeletonisation (**f**). The value of the Dice Similarity Coefficient ($F_1$, asterisk), and $F_{\beta 2}$ statistic (open circle) are shown on the P-R plots and above the corresponding image. The performance of the CNN method remains consistent across the full range of clip-limit tested.

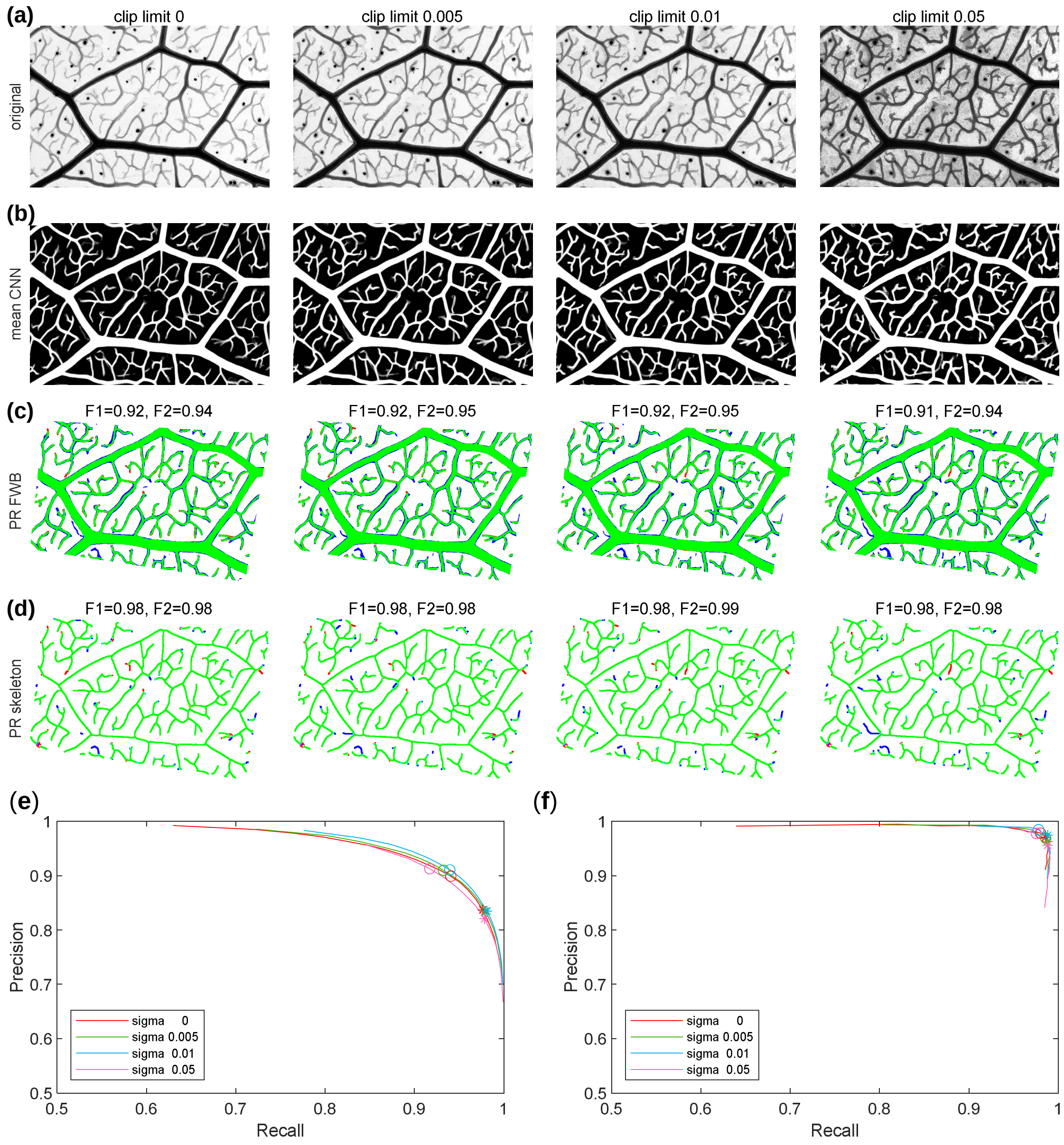

**Fig. S6 Effect of increasing blur on ensemble CNN performance.** A small (~3mm x 2mm) section of a leaf image was blurred using a Gaussian filter with varying radius sigma (**a**). The resultant images were enhanced using an ensemble of three CNNs to give a vein prediction (**b**). The mean CNN predictions were thresholded to give a full-width binary (FWB) image and evaluated against a manually-defined FWB-GT image (**c**), or the resultant single-pixel GT-skeleton (**d**). In each case, true positives ($T_P$) are coded in green, false negatives ($F_N$) in red, and false positives ($F_P$) in blue. Precision-Recall (P-R) plots illustrate the performance of each method as the segmentation threshold is varied for the FWB comparison (**e**) and following skeletonisation (**f**). The value of the Dice Similarity Coefficient ($F_1$, asterisk), and $F_{\beta 2}$ statistic (open circle) are shown on the P-R plots and above the corresponding image. The response of the CNN shows some sensitivity to the level of blur applied.

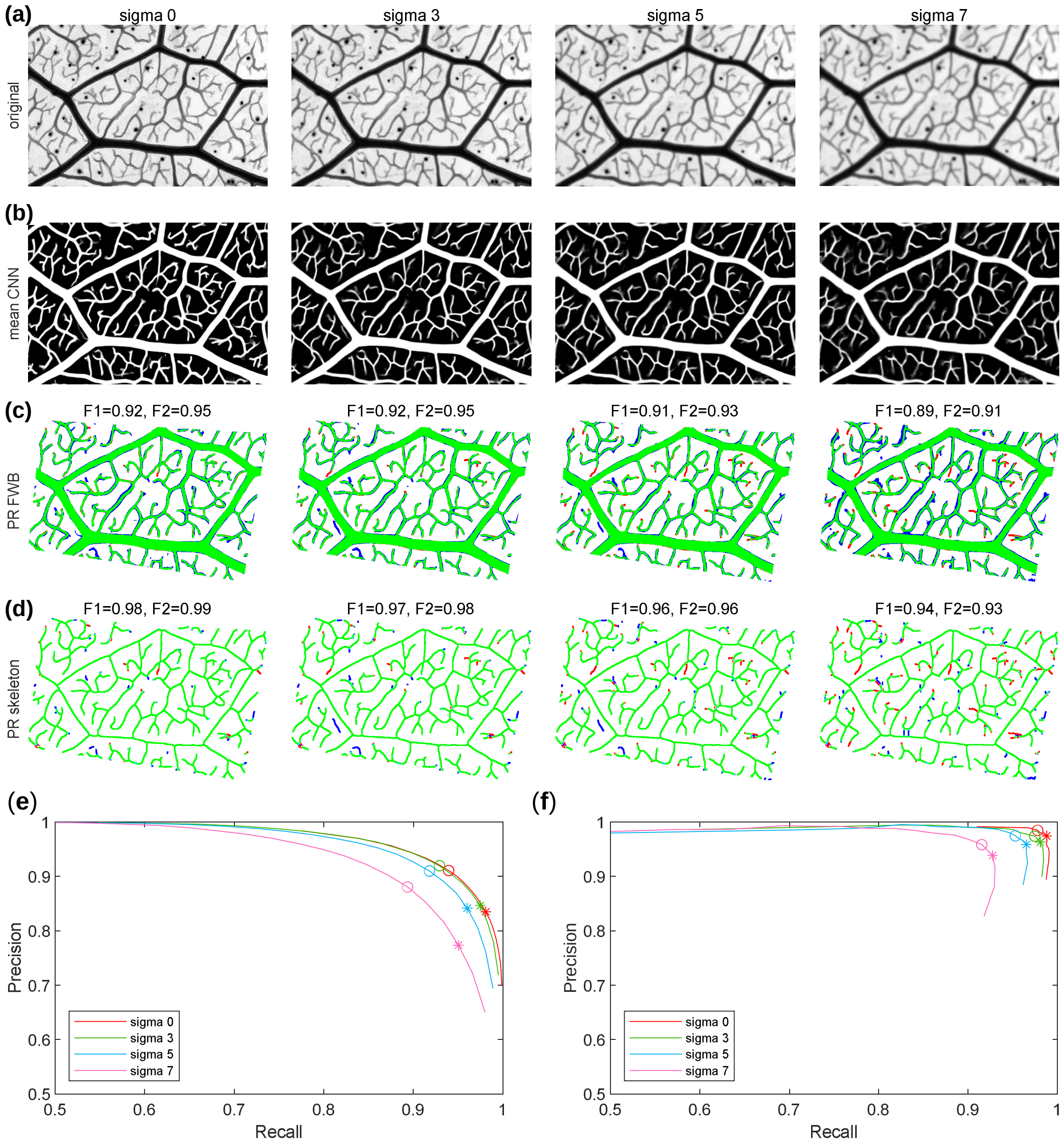

**Fig. S7 Effect of image resolution on ensemble CNN performance.** A small (~3mm x 2mm) section of a leaf image was re-sized to simulate the effect of collecting images at lower spatial resolution (**a**). A scale of 1 corresponds to a pixel size of 1.68 µm, whilst a scale of 0.25 corresponds to 6.72 µm. The resultant images were enhanced using an ensemble of three CNNs to give a vein prediction (**b**). The mean CNN predictions were thresholded to give a full-width binary (FWB) image and evaluated against a manually-defined FWB-GT image (**c**), or the resultant single-pixel GT-skeleton (**d**). In each case, true positives ($T_P$) are coded in green, false negatives ($F_N$) in red, and false positives ($F_P$) in blue. Precision-Recall (P-R) plots illustrate the performance of each method as the segmentation threshold is varied for the FWB comparison (**e**) and following skeletonisation (**f**). The value of the Dice Similarity Coefficient ($F_1$, asterisk), and $F_{\beta 2}$ statistic (open circle) are shown on the P-R plots and above the corresponding image. The response of the CNN is markedly affected by the initial image resolution, failing to capture the finest veins properly as the resolution decreases.

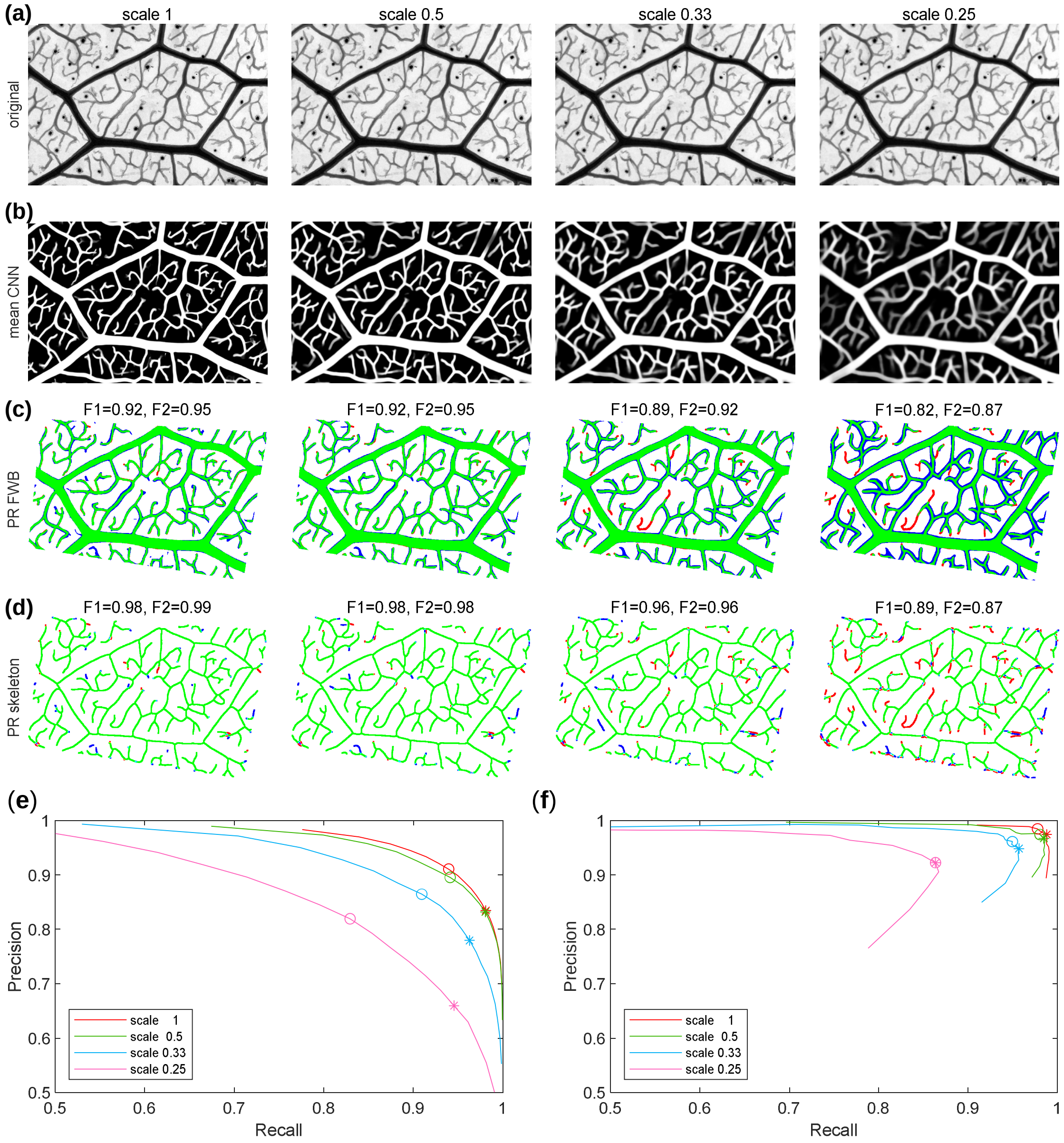

**Fig. S8 Effect of up-sampling to correct for low resolution input images.** A small (~3mm x 2mm) section of a leaf image was re-sized to simulate the effect of collecting images at lower spatial resolution, and then up-sampled again using interpolation to restore the original image size (**a**). The resultant images were enhanced using an ensemble of three CNNs to give a vein prediction (**b**). The mean CNN predictions were thresholded to give a full-width binary (FWB) image and evaluated against a manually-defined FWB-GT image (**c**), or the resultant single-pixel GT-skeleton (**d**). In each case, true positives ($T_P$) are coded in green, false negatives ($F_N$) in red, and false positives ($F_P$) in blue. Precision-Recall (P-R) plots illustrate the performance of each method as the segmentation threshold is varied for the FWB comparison (**e**) and following skeletonisation (**f**). The value of the Dice Similarity Coefficient ($F_1$, asterisk), and $F_{\beta 2}$ statistic (open circle) are shown on the P-R plots and above the corresponding image. Up-sampling is sufficient to almost completely restore the response of the CNN,despite the loss of information in the original lower resolution images.

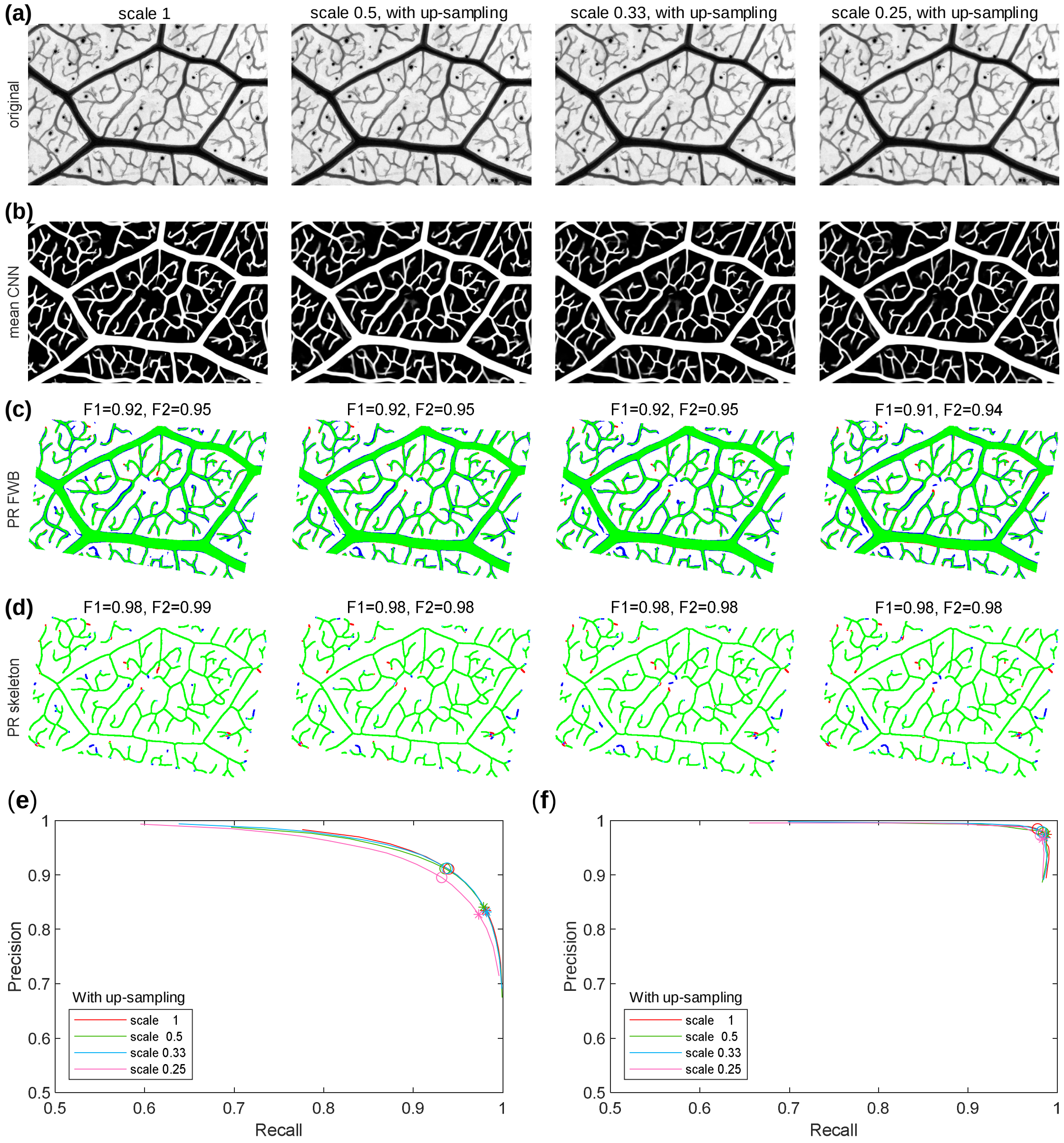

**Fig. S9 Application of LeafVeinCNN to an intact leaf**. (**a**) Original image of a typical intact cleared leaf sample, with inset showing the central region at 3x zoom. Scale bar = 5 mm. (**b**) Average ensemble CNN probability map. (**c**) Pixel-skeleton with loops in magenta and trees in green. (**d**) Pseudo-colour coded vein width skeleton superimposed on the original image. (**e**) Areoles, pseudo-colour coded by $\log_{10}$ area. (**f**) Dual graph connecting centroids of adjacent areoles with the edge weight given by the average width of the intervening vein. The leaf is from is *Quercus turbinella*, collected at the Desert Botanical Garden in Phoenix, Arizona, with the image prepared by Dr. Luiza Aparecido, Miguel Duarte, Sabrina Woo, and Nathan Nguyen

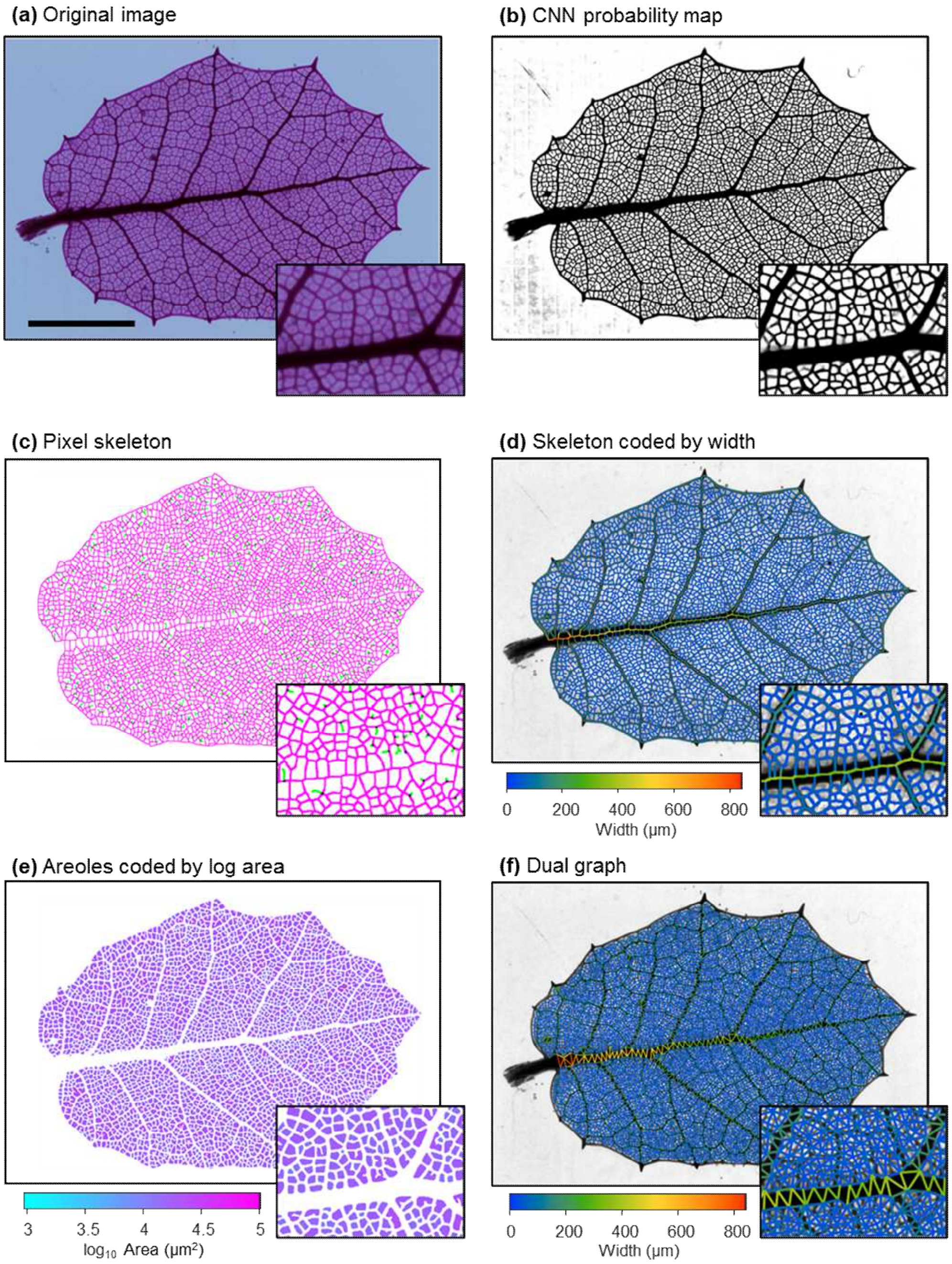

**Table S1 Image enhancement processing parameters**

| Method | Parameter | Value | Output |
|---|---|---|---|
| **All** | Downsample | 2 | |
| **Midgrey** | radius | 45 | Full-width binary image |
| | threshold | [0:0.05:1] | |
| **Niblack** | radius | 45 | Full-width binary image |
| (Niblack, 1985) | threshold | [-0.5:0.05:0.5] | |
| **Bernsen** | radius | 15 | Full-width binary image |
| (Bernsen, 1986) | threshold | [0:0.05:1] | |
| **Sauvola** | radius | 45 | Full-width binary image |
| (Sauvola & Pietikäinen, 2000) | threshold | [0:0.05:1] | |
| **Vesselness** | *fibermetric.m* | | |
| (Frangi *et al.*, 1998) | Image pyramid | 4 levels | Maximum projection of image pyramid |
| | Thickness | 4-9 | |
| | StructureSensitivity | 0.5 * maxhessiannorm | |
| | ObjectPolarity | ‘bright’ | |
| **Phase Congruency** | *Phasecong3.m* | | |
| (Kovesi, 1999; Kovesi, 2000) | Image Pyramid | 2 levels | Maximum projection of image pyramid for the ‘FeatureType’ rescaled between [0 1] |
| | nscale | 5 | |
| | norient | 6 | |
| | minWavelength | 4 | |
| | mult | 2.1 | |
| | sigmaOnf | 0.55 | |
| | k | 2.0 | |
| | cutOff | 0.5 | |
| | g | 10 | |
| | noiseMethod | -1 | |
| **BowlerHat** | *Granulo2D.m* | | |

| | | | |
|---|---|---|---|
| (Sazak *et al.*, 2018) | Image pyramid | 3 levels | Maximum projection of image pyramid |
| | disk size | 2:27 | |
| | orientation | 15 degrees | |
| **MFATλ** | FractionalIstropicTensor.*m* | | Maximum projection of image pyramid |
| (Alhasson *et al.*, 2018) | Image pyramid | 3 levels | |
| | sigmas | [1:1:3] | |
| | spacing | 1 | |
| | whiteondark | true | |
| | tau | 0.2 | |
| | tau2 | 0.5 | |
| | D | 0.01 | |
| **MFATp** | *ProbabiliticFractionalIstropicTensor.m* | | Maximum projection of image pyramid |
| (Alhasson *et al.*, 2018) | Image pyramid | 3 levels | |
| | sigmas | [1:1:3] | |
| | spacing | 1 | |
| | whiteondark | true | |
| | tau | 0.05 | |
| | tau2 | 0.5 | |
| | D | 0.01 | |

**Table S2 Vein metrics**

| **Metric** | **Explanatory notes** | **Units** |
|---|---|---|
| name | The filename | - |
| EndNodes 1 and 2 | Refers to the node ID for node textit{i and node textit{j joined by the edge | - |
| node_Idx 1 and 2 | Refers to the linear index of the node internally in the program, this means that the node pixel can be found from the linear index into the image, as opposed to a position defined by the *x* and *y* co-ordinates | - |
| vein_ID | Each vein gets a specific name (ID) | - |
| Type | Refers to the type of vein (*EL* for Edge Loop, *ET* for Edge Tree, *EB* for Edge Boundary, and *F* for feature = Petiole) | - |

| | | |
|---|---|---|
| Weight | An internal value used for calculations such as the shortest path when it is set to the corresponding parameter | - |
| Width_initial | The average width of the entire vein segment, including the overlap region | pixels |
| Length_initial | The length of the entire vein segment, including the overlap region | pixels |
| Orientation_initial | The orientation of the entire vein segment | radians |
| N pix | The number of pixels in edge | - |
| M pix | The linear index of the middle pixel of the edge | - |
| Intensity and Intensity_cv | Average value and coefficient of variation (CV) for the vein intensity from the original image | arbitrary units |
| Probability and Probability_cv | Average value and CV for the prediction from the CNN | - |
| Length | Length of the vein segment excluding the overlap region | pixels |
| Width and Width_cv | Average width and CV of the vein segment excluding the overlap region | pixels |
| Or_ij and Or_ji | Orientation of the edge between nodes $ij$ or $ij$ measured from the end nodes to the midpoint | degrees |
| Tortuosity | The length of the edge divided by the Euclidean distance between the nodes. Varies from 1 (a straight line) to >1 as the vein is more wiggly | dimensionless |
| Area | Cross-sectional area | pixels$^2$ |
| SurfaceArea | Surface area, excluding overlap region | pixels$^2$ |
| Volume | Volume, excluding overlap region | pixels$^3$ |
| Resistance_2 | Predicted resistance to flow based on the vein area i.e. $l/r^2$ (effectively treats the resistance as scaling with the number of equal sized vessels in a bundle | pixels$^{-1}$ |
| Resistance_4 | Predicted resistance to flow based on Poiseuille flow i.e. $l/r^4$ | pixels-$^{-3}$ |
| Distance | The average Euclidean distance between the vein and the petiole node (if set) | pixels |
| Geodesic | The average geodesic distance along the pixel skeleton between the vein and the petiole node (if set) | pixels |
| Route_factor | The ratio of the geodesic distance to the Euclidean distance | dimensionless |

| | | |
|---|---|---|
| Accessibility | The reciprocal of the average resistance of the shortest path route between node *i* and node *j* to the petiole node (if set) | pixels |
| Ai and Aj | Identifiers of the polygonal areas on either side of the edge | - |

**Table S3 Node metrics**

| Metric | Explanatory notes | Units |
|---|---|---|
| name | The filename | - |
| node_ID | The unique node index | - |
| node_Idx | The linear index into the image for each node | - |
| node_Type | Sets whether the node belongs to an edge, the petiole or is perimeter node connected to the petiole | - |
| node_X_pix and node_Y_pix | Pixel co-ordinates of node | - |
| node_Degree | Number of edges at each node | - |
| node_D1, node_D1 and node_D2 | The diameter of the veins incident at the node, running from largest $D_0$ to smallest $D_2$ | pixels |
| node_Strength | The sum of the widths of the veins at each node | pixels |
| node_Average | Average width of edges at each node | pixels |
| node_OD1, node_OD1 and node_OD2 | Orientation of each vein incident at the node, running from largest $OD_0$ to smallest $OD_2$ | degrees |
| node_AreaRatio | Sum of daughter vein area over parent area $D_1 + D_2/D_0$ | dimensionless |
| node_Symmetry | Ratio of the two daughter veins $D_2/D_1$ | dimensionless |
| theta_D2_D0, theta_D1_D0 and theta_D2_D1 | Branch angles between each pair of veins at a three-way junction | degrees |
| node_Distance | Euclidean distance from the node to the reference point in the petiole | pixels |
| node_Geodesic | Geodesic distance along the skeleton from the node to the reference point in the petiole | pixels |
| node_Route_factor | Ratio of the geodesic distance to the Euclidean distance from the petiole node (if set) | dimensionless |

| | | |
|---|---|---|
| node_Accessibility | The reciprocal of the shortest path route based on resistance between the mid-point of the vein and the petiole node (if set) | pixels |

**Table S4 Areole metrics**

| Metric | Explanatory notes | Units |
|---|---|---|
| name | The filename | - |
| areole_ID | The unique ID for each areole. This corresponds to $A_i$ or $A_j$ in the vein metrics | - |
| node_X_pix and node_Y_pix | The pixel co-ordinates of the areole centroid | pixels |
| Area | The area of the areole | $pixels^2$ |
| MajorAxisLength | The length of the major axis of the ellipse that has the same normalized second central moments as the areole | pixels |
| MinorAxisLength | The length of the minor axis of the ellipse that has the same normalized second central moments as the areole | pixels |
| Eccentricity | The eccentricity is the ratio of the distance between the foci of an ellipse, with the same second moment as the area of the areole, and its major axis length. The value is between 0 and 1, representing a circle and a line, respectively | dimensionless |
| Orientation | The angle between the x-axis and the major axis of the ellipse that has the same second-moments as the areole. The value ranges from -90 to 90 degrees | degrees |
| ConvexArea | The area of the convex hull enclosing the areole | $pixels^2$ |
| Solidity | The proportion of the pixels in the convex hull that are also in the areole. Computed as Area/ConvexArea | dimensionless |
| Perimeter | The distance around the boundary of the areole between each adjoining pair of pixels around the border | pixels |
| MaxDistance | The maximum distance from any pixel within the areole to the bounding veins | pixels |
| MeanDistance | The average distance from all pixels within the areole to the bounding veins | pixels |
| Circularity | The ratio of the perimeter of a circle with the same area to the actual perimeter | dimensionless |
| Elongation | The ratio of the major and minor axis lengths | dimensionless |

| | | |
|---|---|---|
| Roughness | The ratio of the perimeter squared to the area | dimensionless |

**Table S5 Polygon metrics**

| Metric | Explanatory notes | Units |
|---|---|---|
| name | The filename | - |
| polygon_ID | The unique ID for each polygonal area. This corresponds to $A_i$ or $A_j$ in the vein metrics | - |
| node_X_pix and node_Y_pix | The pixel co-ordinates of the centroid of the polygonal area | pixels |
| Area | The area of the polygonal area | pixels$^2$ |
| MajorAxisLength | The length of the major axis of the ellipse that has the same normalized second central moments as the polygonal area | pixels |
| MinorAxisLength | The length of the minor axis of the ellipse that has the same normalized second central moments as the polygonal area | pixels |
| Eccentricity | The eccentricity is the ratio of the distance between the foci of an ellipse, with the same second moment as the area of the polygon, and its major axis length. The value is between 0 and 1, representing a circle and a line, respectively | dimensionless |
| Orientation | The angle between the x-axis and the major axis of the ellipse that has the same second-moments as the polygonal area. The value ranges from -90 to 90 degrees | degrees |
| ConvexArea | The area of the convex hull enclosing the polygonal area | pixels$^2$ |
| Solidity | The proportion of the pixels in the convex hull that are also in the polygonal area. Computed as Area/ConvexArea | dimensionless |
| Perimeter | The distance around the boundary of the polygonal area between each adjoining pair of pixels around the border | pixels |
| MaxDistance | The maximum distance from any pixel within the polygonal area to the bounding veins | pixels |
| MeanDistance | The average distance from all pixels within the polygonal area to the bounding veins | pixels |

| | | |
|---|---|---|
| Circularity | The ratio of the perimeter of a circle with the same area to the actual perimeter | dimensionless |
| Elongation | The ratio of the major and minor axis lengths | dimensionless |
| Roughness | The ratio of the perimeter squared to the area | dimensionless |

**Table S6 HLD metrics**

| Metric | Explanatory notes | Units |
|---|---|---|
| name | The filename | - |
| node_ID | The ID of each node in the dual-graph. During the HLD, new nodes are added that combine the polygonal areas that are fused | - |
| width_threshold | The width of the intervening vein removed during each fusion event | pixels |
| node_Area | The area of the polygons arising from each fusion event | $pixels^2$ |
| node_Degree | The number of the original polygons that have fused prior to each node in the tree | - |
| degree_Asymmetry | The weighted mean value of the asymmetry of the node degrees | - |
| area_Asymmetry | The weighted mean value of the asymmetry of the node areas | - |
| node_HS | Horton-Strahler number | - |
| Perimeter | The perimeter of the new fused polygon | pixels |
| MajorAxisLength | The Major axis length of the new fused polygon | pixels |
| MinorAxisLength | The Minor axis length of the new fused polygon | pixels |
| Eccentricity | The Eccentricity of the new fused polygon | dimensionless |
| Orientation | The Orientation of the new fused polygon | degrees |
| Circularity | The Circularity of the new fused polygon | dimensionless |
| Elongation | The Elongation of the new fused polygon | dimensionless |
| Roughness | The Roughness of the new fused polygon | dimensionless |
| VTotLen | The Total length of the remaining network after vein removal | pixels |
| VTotVol | The Total volume of the remaining network after vein removal | $pixels^3$ |
| MSTRatio | The ratio of the MST for the network divided by the total length | dimensionless |
| Boundary | Flag to indicate whether the polygon is touching the boundary | - |

**Table S7 Summary statistics for vein, node, areole and polygon metrics**

| Metric | Explanatory notes | Units |
|---|---|---|
| **File information** | | |
| File | Filename | - |
| TimeStamp | Time the file was processed | - |
| MicronPerPixel | Calibration value manually entered as a single value for all images in the batch program | $\mu m\ pix^{-1}$ |
| DownSample | Subsampling used to increase processing speed | - |
| **Global Metrics** | | |
| VNE | Veins - Number of Edges | - |
| VNN | Veins - Number of Nodes | - |
| VFE | Veins – Number of Free Ends | - |
| VFEratio | Ratio of Vein Free Ends to total Nodes | dimensionless |
| Valpha | Veins - alpha index or meshedness coefficient (fraction of edges present compared to a maximally connected network) | dimensionless |
| VTotL | Veins - Total Length (total refers to the complete network of both loops and tree regions) | mm |
| VLoopL | Veins - Loop Length (i.e. excludes branching trees leading to free ends) | mm |
| VTreeL | Veins - Tree Length (i.e. length excluding loops) | mm |
| VTotV | Veins - Total Volume (modelled as cylinders from the length and average width. The length excludes the region of overlap with a larger vein) | $mm^3$ |
| VLoopV | Veins - Loop Volume | $mm^3$ |
| VTreeV | Veins - Tree Volume | $mm^3$ |
| VTotW | Average Width for the whole vein network | mm |
| VLoopW | Average Width for the loopy vein network | mm |
| VTreeW | Average Width for the tree vein network | mm |
| VMSTratio | Ratio of the Minimum Spanning Tree connecting all nodes to the total length | dimensionless |
| TotA | Total area included in the analysis (excludes regions that were masked) | $mm^2$ |
| VTotLD | Veins - Total Length Density (total Length over total Area) | $mm^{-1}$ |
| VLoopLD | Veins - Loop Length Density | $mm^{-1}$ |
| VTreeLD | Veins - Tree Length Density | $mm^{-1}$ |
| VTotVD | Veins - Total Volume Density | mm |
| VLoopVD | Veins - Loop Volume Density | mm |
| VTreeVD | Veins - Tree Volume Density | mm |
| VNND | Veins - Node Density | $mm^{-2}$ |
| **Vein metrics** | | |

| | | |
|---|---|---|
| VTot##Len | Veins - Total Length metrics where ## is defined as follows: av = mean; md = median; mo = mode; sd = standard deviation | mm |
| VLoop##Len | Veins - Loop Length metrics | mm |
| VTree##Len | Veins - Tree Length | mm |
| V****##Wid | Average Width, excluding overlap regions of small veins with larger veins, where **** can be Tot, Loop or Tree | mm |
| V-----SAr | Surface area, modelled as a cylinder from the length and average width, excluding overlap | $mm^2$ |
| V-----Vol | Volume modelled as cylinders from the length and width, excluding overlaps | $mm^3$ |
| V-----Tor | Tortuosity – Length of vein / Euclidean distance between end-points | dimensionless |
| V-----Ori | Average orientation of all veins, calculated using circular statistics | degrees |
| **Node metrics** | | |
| NavBD0 | Node average Branch $D_0$. Calculates the average width of the parent branch joining a junction, assuming junctions have degree 3. Junctions with higher degree are very rare. Free end veins have a degree of 1, so only the maximum vein width is recorded | mm |
| NavBD1 | Node average Branch $D_1$. Calculates the average diameter of the largest daughter vein at a junction | mm |
| NavBD2 | Node average Branch $D_2$. Calculates the average diameter of the smallest daughter branch at a junction | mm |
| NavA10 | Node average Angle between $D_1$ and $D_0$. veins at each junction | degrees |
| NavA20 | Node average Angle between $D_2$ and $D_0$. veins at each junction | degrees |
| NavA21 | Node average Angle between $D_2$ and $D_1$ veins at each junction | degrees |
| **Areole metrics** | | |
| ATA | Areole Total Area (Areoles exclude the veins) | $mm^2$ |
| ANN | Areole Number of Nodes | - |
| Aloop | Areole Number of Loops (i.e. the number of Areoles) | - |
| AavAre | Areole average Area | $mm^2$ |
| A---CnA | Areole Convex Area | $mm^2$ |
| A---Ecc | Areole Eccentricity | dimensionless |
| A---Maj | Areole Major axis length | mm |
| A---Min | Areole Minor axis length | mm |
| A---EqD | Areole Equivalent Diameter | mm |
| A---Per | Areole Perimeter | mm |

| | | |
|---|---|---|
| A---Sld | Areole Solidity | dimensionless |
| A---Elg | Areole Elongation | dimensionless |
| A---Cir | Areole Circularity | dimensionless |
| A---Rgh | Areole Roughness | dimensionless |
| A---Ori | Areole Orientation | degrees |
| A---Dav | Areole average Distance to the vein network | mm |
| A---Dmx | Areole maximum Distance to the vein network | mm |
| **Polygon metrics** | | |
| PTA | Polygon Total Area (Polygons include the area up to the pixel skeleton rather than just the areole) | $mm^2$ |
| PNN | Polygon Number of Nodes (excludes those that are incomplete and touching the boundary or internally masked areas) | - |
| Ploop | The number of Polygons | - |
| PavAre | Polygon average Area | $mm^2$ |
| P---CnA | Polygon Convex Area | $mm^2$ |
| P---Ecc | Polygon Eccentricity | dimensionless |
| P---Maj | Polygon Major axis length | mm |
| P---Min | Polygon Minor axis length | mm |
| P---EqD | Polygon Equivalent Diameter | mm |
| P---Per | Polygon Perimeter | mm |
| P---Sld | Polygon Solidity | dimensionless |
| P---Elg | Polygon Elongation | dimensionless |
| P---Cir | Polygon Circularity | dimensionless |
| P---Rgh | Polygon Roughness | dimensionless |
| P---Ori | Polygon Orientation | degrees |
| P---Dav | Polygon average Distance to the vein network | mm |
| P---Dmx | Polygon maximum Distance to the vein network | mm |
